# Incorporating histone H2B variants into chromatin modifies chromatin accessibility to induce epithelial-to-mesenchymal transition in breast cancer

**DOI:** 10.1101/2024.12.21.627414

**Authors:** Hejer Dhahri, Kin H. Lau, Wesley N. Saintilnord, Elisson Lopes, Hannah N. Damico, Flavio R. Palma, Daniël P. Melters, Darrell P. Chandler, Yamini Dalal, Jonathan D. Licht, Marcello G. Bonini, Yvonne Fondufe-Mittendorf

**Affiliations:** Department of Epigenetics, Van Andel Institute, Grand Rapids, MI, 49503; Department of Cellular and Molecular Biochemistry, University of Kentucky, Lexington, KY, 40536; Department of Genetics, Washington University School of Medicine, ST. Louis, MO, 63110; The Edison Family Center of Genome Sciences and Systems Biology, Washington University School of Medicine, St. Louis, MO, 63110; Department of Medicine, Division of Hematology Oncology, Northwestern University Feinberg School of Medicine and the Robert H. Lurie Comprehensive Cancer Center of Chicago, Chicago, IL 60611; Laboratory of Receptor Biology and Gene Expression, National Cancer Institute, NIH, Bethesda, MD, 20894; Division of Hematology/Oncology, University of Florida Health Cancer Center, Gainesville, FL, 32610

## Abstract

Histones scaffold genomic DNA and regulate access to the transcriptional machinery. However, naturally occurring histone variants can alter histone-DNA interactions, DNA and histone modifications, and the chromatin interactome. Hence, alterations in histone variant deposition can disrupt chromatin, and are increasingly recognized as a way to trigger various disease, including cancer. While significant attention has been placed on the biochemical and functional roles of H2A, H3, and H4 histone variants, the variants of H2B remain largely understudied. Here, we show that H2B variants are dysregulated in breast cancer and that certain variants are associated with specific breast cancer subtypes. *HIST1H2BO* overexpression (in particular) is more common in Asian, African American/Black, and young female populations and is associated with a worse prognosis. *In vitro* studies show that H2B1O compacts nucleosome structure. Incorporating H2B1O into chromatin activates pro-inflammatory and oncogenic pathways, induces the epithelial-to-mesenchymal transition (EMT), and generates resistance to first-line chemotherapeutic agents. Thus, H2B1O acts much like an onco-histone, with H2B variant expression being a prognostic biomarker for breast cancer and a potential new target for drug therapies to enhance treatment efficacy.

**Graphical Abstract:** 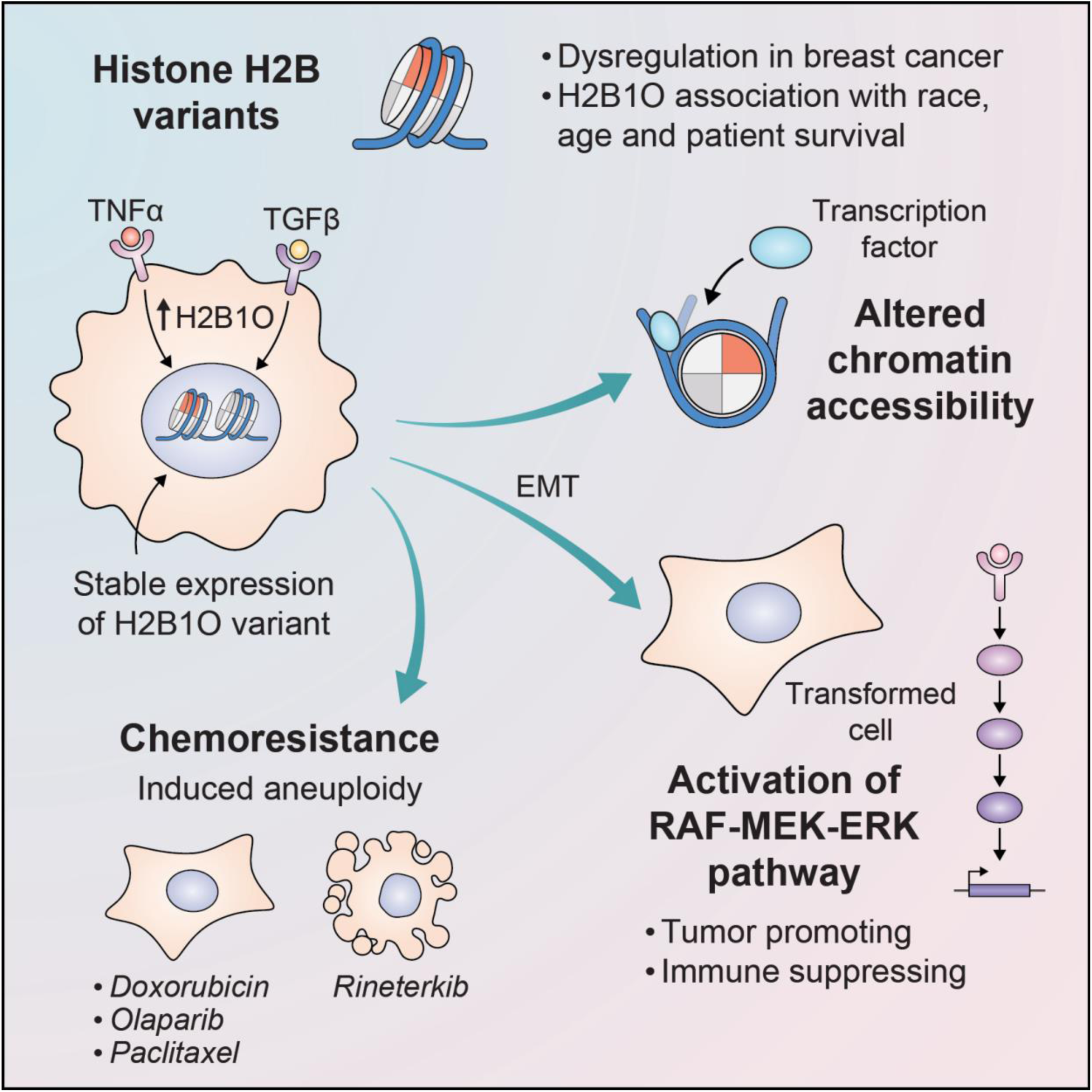

## Introduction

Breast cancer is a complex disease characterized by wide-ranging genetic and epigenetic alterations. Inter-tumor heterogeneity results in diverse clinical presentations across patients, to include significant variations in angiogenic, invasive, and metastatic potential [1–5], and has a significant impact on patient prognosis, treatment options, response to therapy, and metastatic sites [6–8]. Advances in genomic/transcriptomic profiling and histopathological examinations have enabled the field to identify new and distinct factors contributing to breast cancer etiology, such that breast cancer can now be categorized into luminal A, luminal B, human epidermal growth factor receptor 2 positive (HER2+), triple-negative breast cancer (TNBC), and basal-like non-TNBC molecular sub-types. This classification scheme is largely based on the estrogen receptor (ER), progesterone receptor (PR), and HER2 receptor expression levels [9–11], and is widely used to guide treatment decisions [12]. For instance, patients with luminal tumors generally respond to endocrine therapy and occasionally chemotherapy [13, 14]. Tumors that are HER2+ exhibit *ERBB2* oncogene amplification and overexpression [15] and can be managed with specific anti-HER2 treatments, including monoclonal antibodies and tyrosine kinase inhibitors [16–18]. Basal-like tumors that lack hormone receptor and *HER2* amplification are more recalcitrant and typically treated with standard chemotherapy, with only ∼20% of these tumors responding to most FDA-approved therapies [19, 20]. For these patients, the identification of new, more effective treatments remains a top priority.

Currently, several breast cancer epigenetic therapies are being explored, including histone deacetylase inhibitors (HDACi), DNA methyltransferase inhibitors (DNMTi), and lysine methyltransferase inhibitors (KMTi), alone or in combination with immunotherapy [21, 22]. Several of these epigenetic drugs seem to be effective for breast cancer [23, 24]. Nevertheless, breast cancer is among the most heterogeneous malignancies. For instance, there are multiple tumor sub-classes within individual breast cancer subtypes, each with its own molecular features, sensitivity to therapeutic agents, and resistance mechanisms [25, 26]. There is also well-documented intra-tumor heterogeneity, which enables tumors to evolve rapidly in response to environmental cues or treatments, often through epigenetic reprogramming [27–29]. Thus, there is considerable interest in understanding those epigenetic factors that contribute to breast cancer etiology and/or treatment resistance so the field can develop more precise epigenetic-based therapies.

Histones and histone post-translational modifications (PTMs) control how transcriptional machinery accesses and interacts with genomic DNA. Thus, disrupting or dysregulating histone biology can alter gene expression. Histone proteins are usually categorized as canonical or histone variants—often considered isoforms of the canonical histones. In contrast to canonical histones, which are typically deposited into chromatin during the S phase of the cell cycle, histone variants are expressed and incorporated into chromatin throughout different cell cycle phases [30]. Importantly, accumulating evidence suggests that dysregulated incorporation of histone variants into chromatin can contribute to cancer initiation and progression, with most prior studies focused on H2A, H3, and H4 variants. For example, H2A.Z expression is implicated in breast cancer progression and is associated with lymph node metastasis [31–33]. An increased deposition of histone H3.3 into chromatin, accompanied by a reduction in H3.1 levels, was also observed during breast cancer metastasis [34, 35]. Finally, H4G regulates transcription and cell-cycle progression and exhibits a tumor-stage dependent over-expression [36, 37]. Thus, histone variants are involved in breast cancer, but there has been limited research on H2B variants and their role in oncogenesis.

H2B is encoded by several genes, with five genes encoding the same protein (H2B1C) and the others encoding distinct isoforms that differ from H2B1C by one or a few amino acids. We recently showed that histone H2B variants are differentially expressed in multiple cancers [38]. In this study, we therefore investigated how H2B variants are dysregulated in breast cancer. We demonstrate that the majority of H2B genes are dysregulated in breast cancer patients, with *HIST1H2BO*, *HIST1H2BH*, and *HIST1H2BC* (encoding H2B1O, H2B1H, and H2B1C respectively) displaying some of the most pronounced differential expression across and within distinct molecular subtypes. We also show that H2B variants (which vary from each other by only a few amino acids) exhibit differential expression patterns and occupy specific genomic locations. H2B1O dysregulation, in particular, is linked to the most aggressive subtypes of breast cancer, alters chromatin accessibility, activates oncogenic pathways, induces the epithelial-to-mesenchymal transition (EMT), and affects transformed breast epithelial cell response to chemotherapy. H2B1O, therefore acts much like an onco-histone, with prognostic implications for breast cancer.

## Results

### H2B variants are highly dysregulated across cancers, and their expression is specific to tumor type

We previously analyzed patient transcriptome data from The Cancer Genome Atlas (TCGA) to compare histone H2B variant gene expression in various cancers relative to their corresponding normal tissue types [38]. We identified 18 H2B genes with tumor-specific dysregulation, with *HIST1H2BO* showing the highest differential expression in breast cancer (**Fig 1A** and **S1A**). We next used methods outlined in Thennavan et al. [39] to parse TCGA breast cancer data into the five most common molecular subtypes (**Fig S1B**). We again found significant H2B variant gene expression differences in each breast cancer subtype (**Fig S1C**). *HIST1H2BO* exhibited the greatest differential expression among these variants, especially in the most aggressive breast cancer subtypes (i.e., TNBC and HER2-enriched; **Fig 1B**).

**Fig 1.**
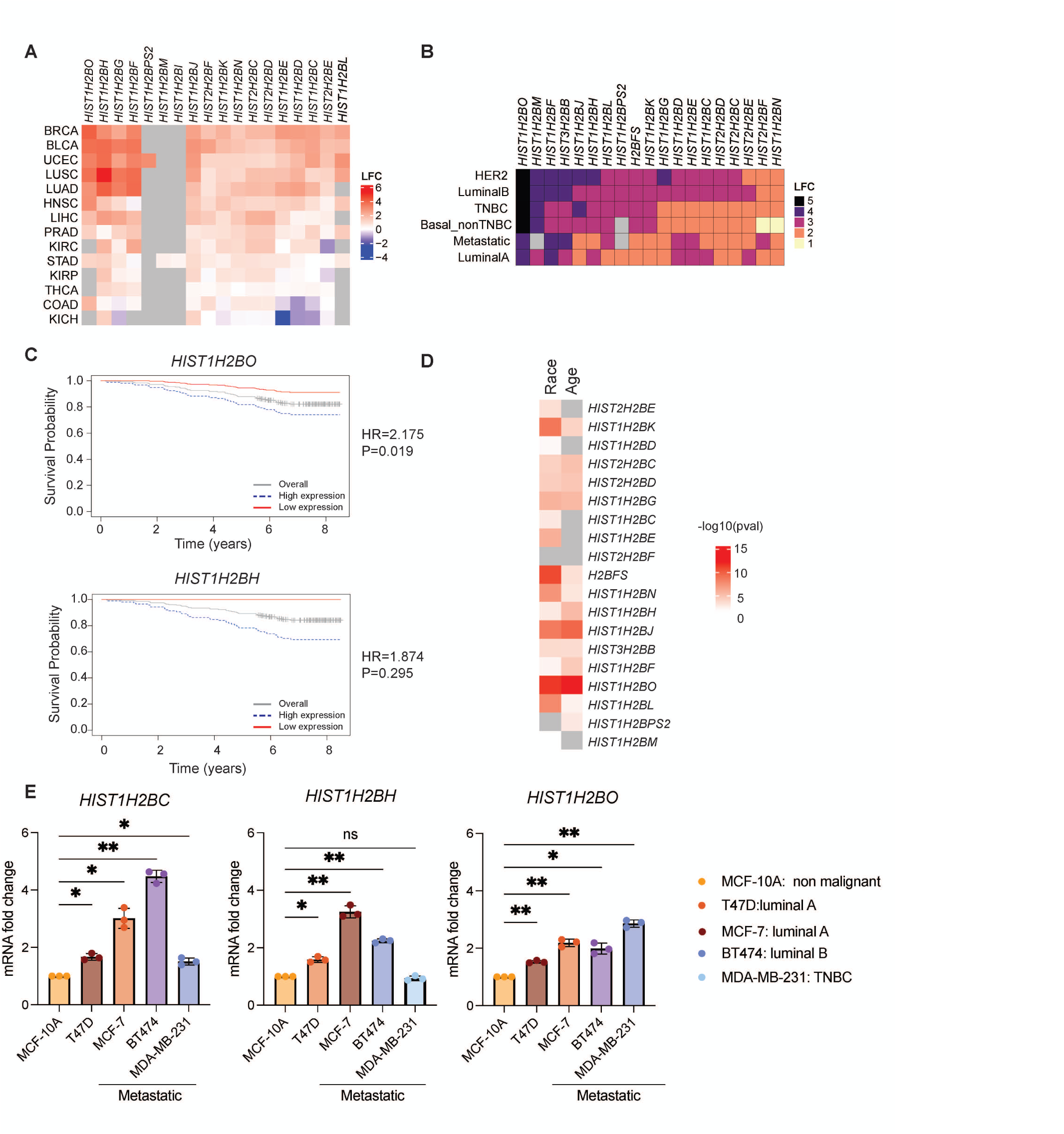
H2B variants are dysregulated in breast cancer. **A**, H2B gene expression in tumors vs. adjacent normal tissues. Data are from the TCGA database, with p_adj_ <0.05. Gray boxes indicate no significant difference in gene expression. LFC = Log_2_(FoldChange). BRCA = breast invasive carcinoma; BLCA = bladder urothelial carcinoma; UCEC = uterine corpus endometrial carcinoma; LUAD = lung adenocarcinoma; LUSC = lung squamous cell carcinoma; LIHC = liver hepatocellular carcinoma; PRAD = prostate adenocarcinoma; KIRC = kidney renal clear cell carcinoma; COAD = colon adenocarcinoma; STAD = stomach adenocarcinoma; KIRP = kidney renal papillary cell carcinoma; THCA = thyroid carcinoma; and KICH = kidney chromophobe. **B**, Differential expression of H2B genes across breast cancer subtypes. **C**, Kaplan–Meier plots of overall survival for patients over-expressing *HIST1H2BO* or *HISTH2BH*. p values as determined by log rank test for all analyses and Hazard ratio (HR) are provided in the graph. **D**, Correlation between H2B variant gene expression, age, and race. **E**, *HIST1H2BC*, *HIST1H2BH*, and *HIST1H2BO* expression in common immortalized breast cancer cell lines relative to the non-cancerous MCF10A cell line.

Next, we used microarray data from the PrognoScan database [40] to determine the prognostic value of H2B variant gene expression. Kaplan-Meier curves revealed that high *HIST1H2BO* and *HIST1H2BH* expression is significantly associated with poor overall survival in breast cancer (**Fig 1C**) and other cancers (**Fig S1D, E**). We also interrogated whether H2B variant expression correlates with age and race since these are well-acknowledged risk factors for aggressive breast cancers [41–43]. We found that *HIST1H2BFS*, *HIST1H2BJ*, *HIST1H2BK*, HIST1H2BL, *HIST1H2BN,* and *HIST1H2BO* gene expression correlates with race; *HIST1H2BJ* and *HIST1H2BO* expression correlates with age; and *HIST1H2BO* has the most significant correlation with both race and age (**Fig 1D**). Interestingly, we also found that *HIST1H2BC*, *HIST1H2BH*, and *HIST1H2BO* expression was significantly elevated in several common breast cancer cell lines compared to the non-malignant epithelial breast cell lines MCF10A, with *HIST1H2BO* having the highest expression in a highly invasive TNBC cell line (**Fig 1E**). Thus, H2B variant expression is associated with breast cancer including invasiveness and aggressiveness, with *HIST1H2BO* expression (in particular) associated with patient demographics and breast cancer aggressiveness.

### *HIST1H2BO* expression heterogeneity in breast cancer

When analyzing histone variant gene expression in breast cancer samples from the TCGA, we noticed considerable heterogeneity in the expression of H2B variants and other chromatin regulators, even among breast cancer samples from the same molecular subtypes (**Fig 2A** and **S2**; n=86 chromatin-related genes). We therefore stratified the TCGA breast cancer data into “high”, “medium”, and “low” expression groups, where each group was defined based on the coordinated expression of these 86 genes. We then used EPIC, TIMER, XCELL, and PUREE to remove non-tumor signals, and conducted gene set enrichment analysis (GSEA) on the bulk transcriptomes from each group to identify potentially dysregulated pathways associated with *HIST1H2BO* and its co-expressed chromatin-regulating genes.

**Fig 2.**
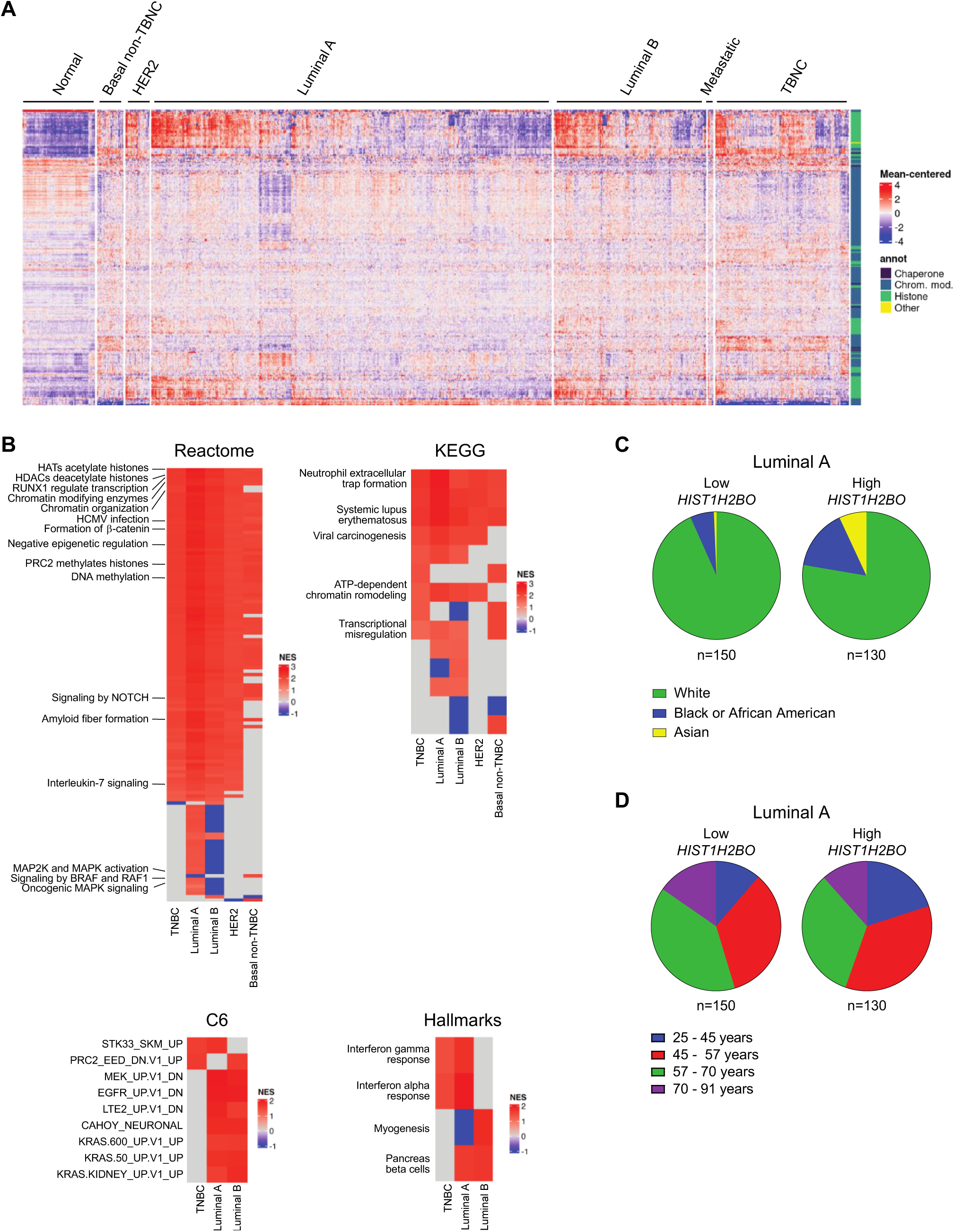
H2B variants and chromatin-modifying genes are differentially expressed within breast cancer subtypes. **A**, Chromatin-modifying gene expression in breast cancer subtypes. Data are from The Cancer Genome Atlas (TCGA) database. Each column represents a patient tumor sample. **B**, Pathway enrichments of differentially expressed genes in breast cancer subtypes with “high” versus “low” *HIST1H2BO* expression. Shared pathways include those associated with inflammation, tumor migration, and oncogenic signatures. **C**, Breast cancer patients with high *HIST1H2BO* expression have more Asian and African Americans than that group of patients with low *HIST1H2BO* expression. **D**, Same as **C**, except patients stratified by age. Patients with high *HIST1H2BO* expression tend to be younger than those with low *HIST1H2BO* expression.

We next compared transcriptomes from the high versus low chromatin-related expression clusters. As expected, those patients with the highest *HIST1H2BO* expression had significantly up-regulated pathways related to chromatin organization, chromatin-modifying enzymes, epigenetic regulation of gene expression, histone deacetylases (HDACs), histone acetyltransferases (HATs), and ATP-dependent chromatin remodeling, regardless of breast cancer subtype (**Fig S2**). Surprisingly, all breast cancer patients with elevated *HIST1H2BO* expression had activated pathways associated with human cytomegalovirus (HCMV) infection, neutrophil extracellular trap formation, systemic lupus erythematosus, and viral carcinogenesis (**Fig 2B**). Within the luminal B subtype, platelet activation, toll like receptor 2 (TLR2), 4 (TLR4), and TLR6:TLR2 cascades were suppressed (**Fig S2**). Patients with high *HIST1H2BO* expression also had significant enrichment of oncogenic pathways associated with poor prognosis (e.g., KRAS signatures; **Fig 2B**, C6 signatures). These data are consistent with the well-established, strong connection between viral infections and tumor formation [44, 45].

Finally, we found that the group of patients with high *HIST1H2BO* expression have more Asian and African Americans relative to the groups with medium or low *HIST1H2BO* expression (**Fig 2C** and **Fig S3A**). Patients with high *HIST1H2BO* expression also had significantly more women between 25-45 years of age (**Fig 2D** and **Fig S3B**). These findings are significant because aggressive breast cancer subtypes (like TNBC) are more prevalent among younger African American and Asian women [46–48]. Our results suggest that *HIST1H2BO* over-expression is linked to oncogenic, pro-inflammatory, and immune-related processes, especially in young African American and Asian women.

### H2B variants alter nucleosome structure

The primary amino acid sequences of H2B1C, H2B1H, and H2B1O differ from each other by only one or two amino acid residues (**Fig 3A**). The histone fold domain consists of a core helix (α2) coupled to two shorter α-helices (α1 and α3) flanked by L1 and L2 loops, respectively. Residue 39 is situated at α1 and facilitates the interaction between histones and nucleosomal DNA. H2B also has an elongated N-terminal tail that extends between two DNA gyres and stabilizes the interaction between H2B and DNA. These interactions are important for chromosome condensation and forming higher-order chromatin structures [49–51]. To determine if these seemingly minor amino acid differences alter nucleosome stability, we reconstituted H2B1C-, H2B1H-, and H2B1O-containing nucleosomes using ^32^P-labeled 601 Widom sequence flanked by a 50 bp linker DNA at both ends.

**Fig 3.**
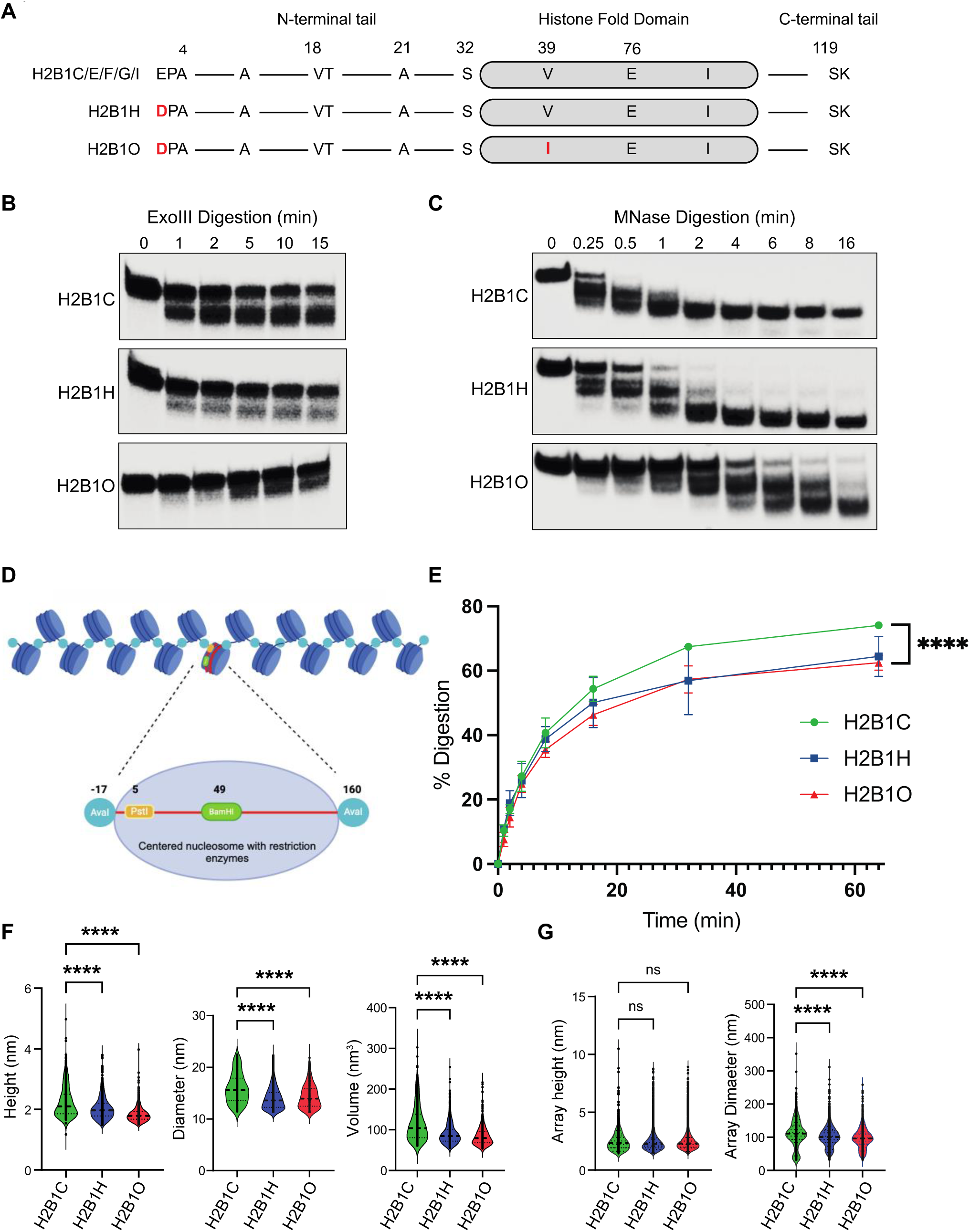
HIST1H2BO alters nucleosome and chromatin structure. **A**, H2B variant sequence alignment. Red letters indicate amino acid substitutions relative to the most common isoforms. **B**, Gel electrophoresis images of H2B variant nucleosomes digested with Exonuclease III for the indicated times. H2B1O nucleosome DNA digests slower, indicating reduced enzyme access to the nucleosome DNA. Each experiment was repeated four times. **C**, Same as in **B**, except with MNase. **D**, 17-mer H2B variant nucleosome arrays with strategically designed restriction enzyme cut sites. **E**, Same as in **B**, except with Pst1. **F**, Nucleosome height, diameter, and volume as quantified by atomic force microscopy. **G**, Nucleosome array height and diameter measured by atomic force microscopy.

All H2B variants efficiently incorporated into and formed stable mono-nucleosomes (**Fig S4A, B**). An exonuclease III digestion assay [52] showed that the linker DNA in H2B1O nucleosomes was the most resistant to enzymatic digestion (**Fig 3B**), suggesting that the linker DNA in H2B1O nucleosomes is less available for enzymatic digestion than in H2B1C or H2B1H nucleosomes. These data were confirmed with an MNase digestion assay [38, 53] (where MNase has endo-and exonuclease activities; **Fig 3C**). We then used a DNase I digestion assay [54] to test if incorporating these H2B variants affected the translational positioning of the nucleosome. As with exonuclease III and MNase digestions, the DNA in H2B1O nucleosomes was less accessible to DNase I than H2B1H or H2B1C nucleosomes (**Fig S4C**). These data suggest that H2B1O strengthens the interaction between the core histone octamer and nucleosome/linker DNA, making nucleosomal DNA less accessible to digestion *in vitro*.

Changes in linker DNA flexibility could influence higher-order chromatin structure. To test this possibility, we reconstituted 17-mer nucleosome arrays with either H2B1C-, H2B1H-, or H2B1O-containing histone octamers, and tandem repeats of the Widom 601 DNA nucleosome positioning sequence (NPS), where the DNA construct contained a BamHI restriction site at the center of the middle nucleosome, AvaI restriction sites at the boundaries of each mono-nucleosome, and PstI restriction sites near the entry and exit site of the middle nucleosome (**Fig 3D**; [38, 55, 56]). All nucleosome arrays were efficiently reconstituted (**Fig S4D**) and there were no detectable nucleosome-free 601 DNA fragments (which would be detected by BamHI and AvaI digestion assays; **Fig S4E**). A *PstI* restriction enzyme assay again showed that the H2B1O (and H2B1H) nucleosome DNA entry/exit sites were less accessible than in H2B1C nucleosome arrays (**Fig 3E, Fig S4F**), which suggests that H2B1O- and H2B1H-containing nucleosome arrays are more compact than H2B1C chromatin. We confirmed these findings by atomic force microscopy (AFM), which creates a topological image of the nucleosomes and arrays at sub-nanometer resolution (**Fig S4G**). Nucleosome height and diameter were smaller in H2B1O- and H2B1H-containing nucleosomes, which translated into smaller volume (**Fig 3F**). For nucleosomal array, there was no significant change in array height, indicating that the nucleosome arrays were efficiently distributed across the atomically flat AFM surface. However, we did see a significant reduction in the maximum Feret’s diameter in H2B1H- and H2B1O-nucleosome arrays (**Fig 3G**). Thus, H2B1O (in particular) and H2B1H generate smaller nucleosomes, more compact chromatin arrays, and decreases enzyme access to linker and nucleosomal DNA, all of which could influence DNA transcription.

### Incorporating H2B1O into chromatin activates oncogenic and inflammatory pathways

To understand how H2B variants regulate gene expression and oncogenesis, we used an experimental flow as depicted in (**Fig 4A**). We first designed a minigene system (**Fig 4B**; [38]) to express either HA-tagged *HIST1H2BC*, *HIST1H2BH*, or *HIST1H2BO* in the non-tumorigenic MCF10A epithelial cell line, where cells transfected with an empty vector (EV) served as controls. Each variant was stably expressed throughout all cell cycle phases (**Fig S5A, B**), and immunoblots showed that each HA-tagged H2B variant was translated and incorporated into chromatin (**Fig 4C** and **S5C**).

**Fig 4.**
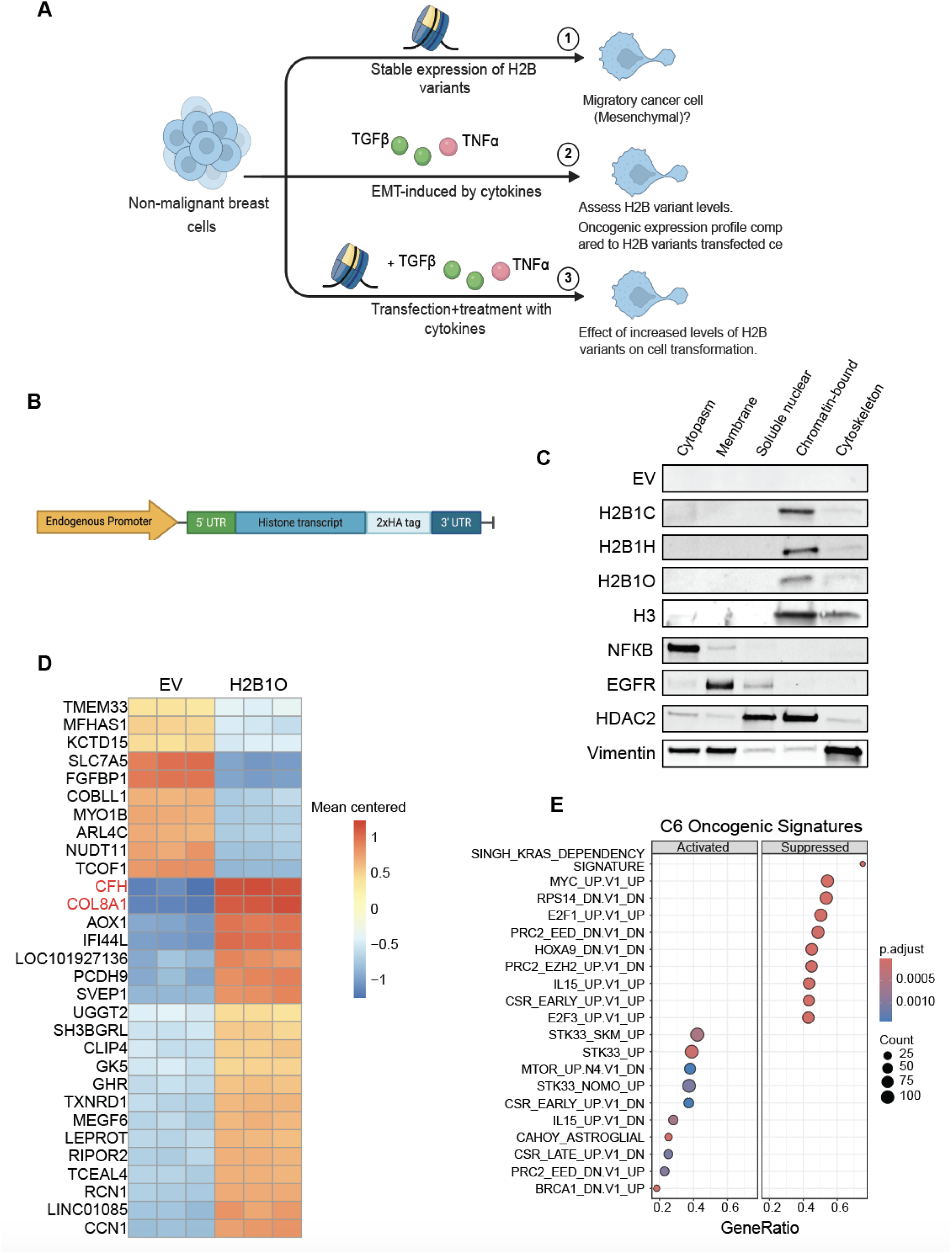
H2B variants activate distinct cellular pathways in non-cancerous breast epithelial cells. A, The experimental design of our in vitro cellular experiments. **B**, Minigene system for stably expressing H2B variants in MCF10A cells. **C**, Immunoblots show that H2B variants co-localize with H3 in the chromatin-bound sub-cellular fraction. EV = empty vector **D**, The top 30 differentially expressed genes (DEGs) in MCF10A cells that stably express *HIST1H2BO*. **E**, The top enriched MSigDB C6 oncogenic signatures in MCF10A cells stably expressing *HIST1H2BO*.

Next, we performed RNA-seq and GSEA to identify differentially expressed genes and pathways in cells that were expressing and incorporating H2B variants into their chromatin. Cells expressing *HIST1H2BO*, in particular, were enriched for several breast cancer-associated genes and pathways (**Fig 4D,E**), including complement factor H (CFH) and COL8A1, both of which are linked to aggressive breast cancer phenotypes and adverse outcomes [57–61]. In addition, these *HIST1H2BO* expressing cells were enriched for breast cancer (e.g., BRCA1_DN.V1_UP; KRAS.LUNG.BREAST_UP.V1_UP; and KRAS.BREAST_UP.V1_UP) and other oncogenic pathways (e.g., STK33_SKM_UP; STK33_UP; and STK33_NOMO_UP), inflammatory pathways (viral mRNA translation, herpes simplex virus 1 infection, and interferon alpha response), and chromatin modification pathways (e.g., activation of PRC2_EED_DN.V1_UP; and suppression of PRC2_EED_DN.V1_DN and PRC2_EZH2_UP.V1_DN; **Figs S6** and **4E**).

Cells expressing *HIST1H2BC* showed increased *TXNRD1* expression, which is associated with an unfavorable breast cancer prognosis [62] (**Fig S7A**). *HIST2H2BC* expression suppressed cell adhesion, extracellular matrix organization, cell junction organization, and cell-cell communication pathways (**Fig S7A**), suggesting that H2B1C may regulate cell migration. On the other hand, *HIST1H2BH*-expressing cells had up-regulated genes linked to immunoregulatory and inflammatory processes (including TNFAIP3, TNF1IP6, CXCL1, and CXCL2; **Fig S7B**) that are associated with the regulation of tumor-associated neutrophils and anti-tumor immune suppression in breast cancer [63–66]. Cells expressing *HIST1H2BH* also had activated pro-inflammatory pathways, including INTERFERON_GRAMMA_RESPONSE, interleukins signaling, TNFA_SIGNALING_VIA_ NFKB, and INFLAMMATORY_RESPONSE (**Fig S7B**). These data suggest that H2B variants have distinct regulatory roles and that H2B1O (in particular) is influencing chromatin structure to activate EMT, inflammatory, and oncogenic pathways.

### H2B1O localizes to the promoter regions of oncogenic and inflammatory genes

To identify the genomic distribution of H2B variants and determine if their incorporation correlates with changes in gene expression, we conducted CUT&RUN analysis followed by an over-representation analysis on differentially expressed genes (DEGs). Interestingly, H2B1O was predominantly localized to promoters of upregulated genes associated with known breast cancer pathways, including STK33_UP, STK33_NOMO_UP, and MTOR_UP. N4.V1_DN (**Fig 5A**). H2B1H was mainly deposited at the promoters of inflammatory response genes, and these same genes/promoters were also enriched for the activating H3K4me3 mark (**Fig 5B**). Interestingly, only cells expressing H2B1O had activating H3K4me3 marks at the promoters of EMT pathway genes (**Fig 5A** and **B**).

**Fig 5.**
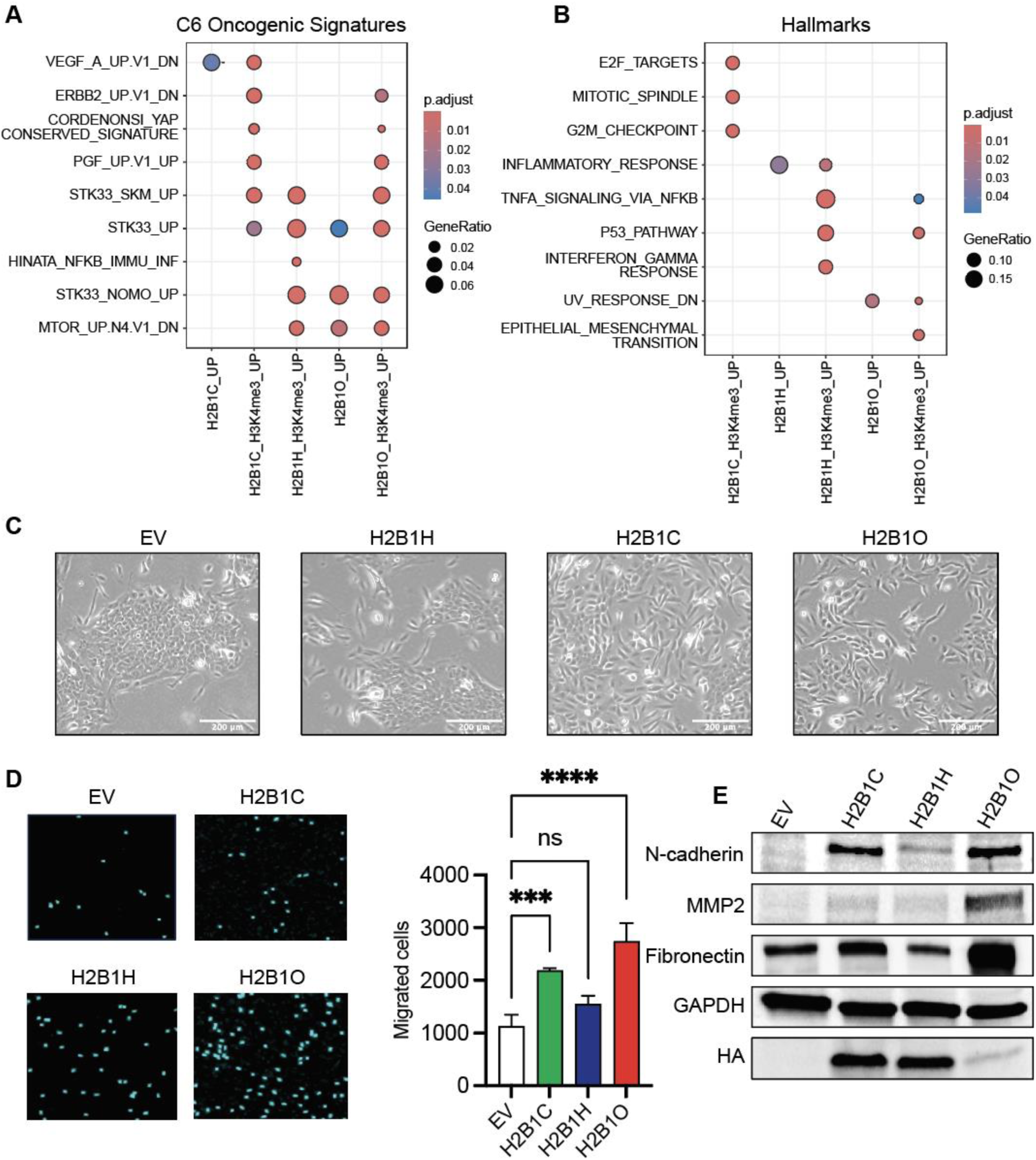
H2B variants occupy oncogene and inflammatory pathway promoters. **A**, CUT&RUN over-representation analysis of C6 oncogenic signatures and activating H3K4me3 marks in MCF10A cells transfected with H2B variant minigenes. Cells transfected with an empty vector (EV) served as controls. **B**, Same as **A**, except for Hallmark signatures. **C**, Microscope images of MCF10A cells stably transfected with H2B variants minigenes. An elongated, spindle-like morphology is indicative of cells undergoing the epithelial-to-mesenchymal transition (EMT). Scale bar is 200 µm. **D**, Migrating cells 24 h after seeding in a transwell insert (n=3; biological replicates); **** p<0.0001. **E**, Immunoblot of HA-tagged H2B variants and EMT markers in MCF10A cells stably transfected with H2B variant minigenes. Cells expressing *HIST1H2BO* expressed the highest level of mesenchymal markers.

Cells transfected with an empty vector retained the cobblestone-like shape characteristic of MCF10A cells, but cells expressing *HIST1H2BC*, *HIST1H2BH*, and *HIST1H2BO* all showed evidence of an elongated, spindle-like morphology typical of mesenchymal cells, a phenotype that was most pronounced in cells expressing *HIST1H2BO* (**Fig 5C**). These results suggest that cells expressing H2B variants are in different stages of the EMT, with most of the H2B1O-transfected cells having transitioned to the mesenchymal state. These results were confirmed with a trans-well migration assay (**Fig 5D**) and are also supported by elevated N-cadherin, matrix metallopeptidase 2 (MMP2), and fibronectin expression (**Fig 5E**; N-cadherin and MMP2 are EMT markers, and fibronectin is involved in cell growth, migration, and metastasis). Thus, H2B1C, H2B1H, and H2B1O are enriched at different chromatin regions, associate with different (yet activated) oncogenic pathways, and generate different cell migration potentials, suggesting that H2B variants (and H2B1O in particular) regulate the EMT.

### EMT-inducing cytokines increase H2B variant expression and trigger an H2B1O-like gene expression profile

To investigate if these H2B variants are important for reprogramming cells into a mesenchymal state, we treated non-transfected MCF10A breast epithelial cells with tumor necrosis factor-alpha (TNF-α) and transforming growth factor β (TGF-β) to drive all cells through the EMT (**Fig 6A**; [67–69]). As expected, there was decreased epithelial marker expression, increased mesenchymal marker expression (**Fig 6B, C**), and up-regulation of several tumor-promoting genes (**Fig S8**). Treated cells also activated several important oncogenesis, inflammation, and tumor migration pathways (**Fig S8**). Interestingly, inducing the EMT with TNF-α and TGF-β increased the endogenous levels of *HIST1H2BC*, *HIST1H2BH*, and *HIST1H2BO* expression, with the most pronounced increase in *HIST1H2BO* expression (**Fig 6D**).

**Fig 6.**
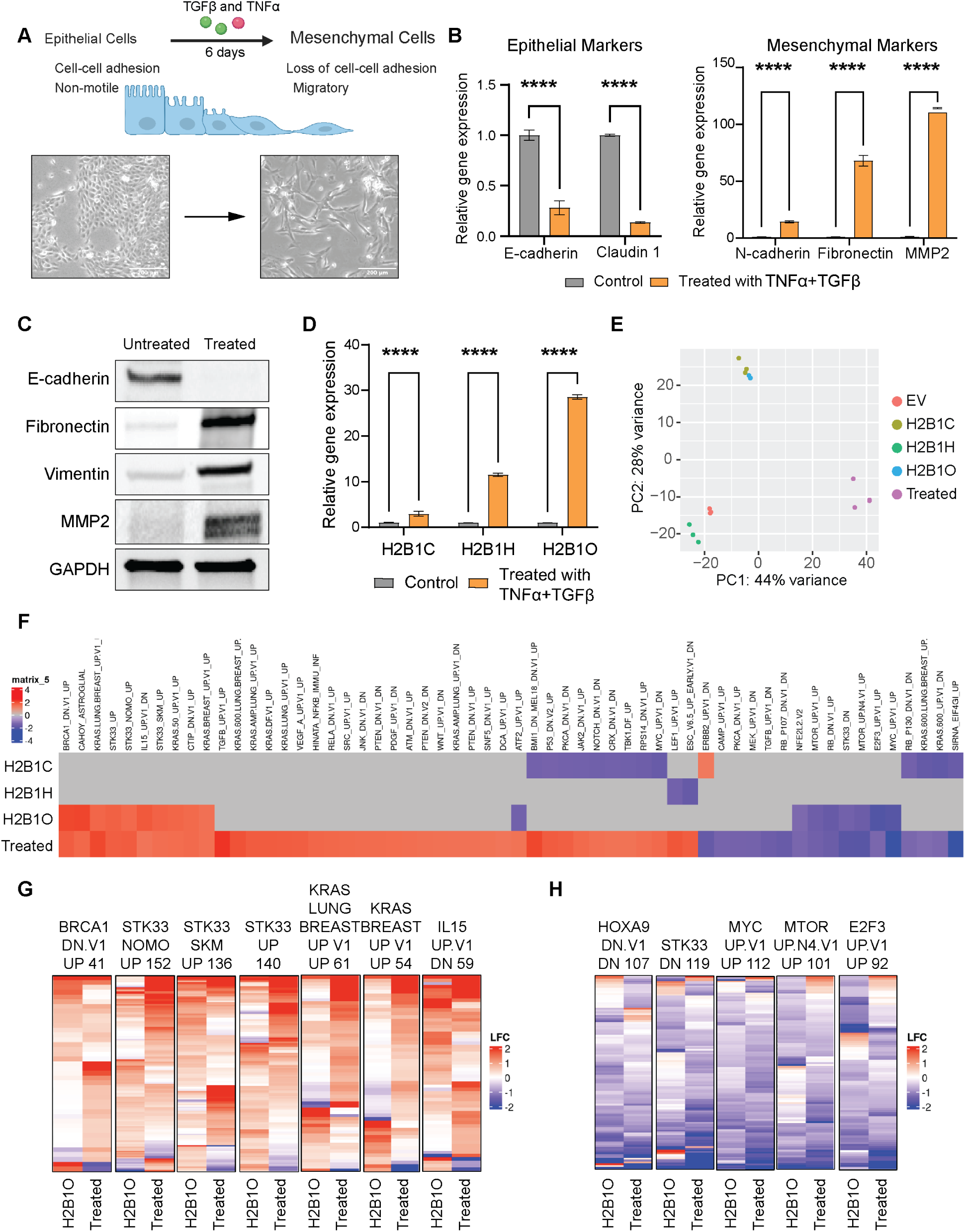
Inducing the epithelial-to-mesenchymal transition (EMT) generates transcriptional responses similar to those induced by HIST1H2BO. **A**, Cytokine-induced transformation of MCF10A cells, and representative microscope images of non-treated and transformed cells. Scale bar is 200 µm. **B**, Relative expression of epithelial and mesenchymal markers in untreated and cytokine-transformed MCF10A cells (n=3); **** p<0.0001. **C**, Immunoblots of epithelial and mesenchymal markers. **D**, Relative gene expression of H2B variants in untreated and cytokine-transformed MCF10A cells (n=3); **** p<0.0001. **E**, PCA plot of total RNA-seq profiles. **F**, Top enriched C6 oncogenic signatures in MCF10A cells stably expressing H2B variant minigenes, and in cytokine-transformed cells. **G-H**, MCF10A cells expressing *HIST1H2BO* have similar oncogenic gene expression patterns as cytokine-transformed cells. Each column is a MSigDB C6 oncogenic signatures; each row is a gene within the respective signature. LFC = log fold-change.

Gene expression profiles from cytokine-transformed cells and cells expressing H2B variants separated into distinct clusters by PCA (**Fig 6E**). Cells expressing *HIST1H2BC* and *HIST1H2BO* clustered together and more closely to cytokine-transformed cells than did cells expressing *HIST1H2BH* (or those containing an empty vector), indicating that they generate gene expression profiles similar to cytokine-induced EMT. Cells expressing *HIST1H2BC* were enriched in MSigDB oncogenic signatures and gene ontology terms, but only cells expressing *HIST1H2BO* showed any overlap with the EMT transformed cells (**Fig 6F**). Within these overlapping oncogenic pathways, cells expressing *HIST1H2BO* had gene expression patterns similar to those in cytokine-transformed cells (**Fig 6G, H**). These data suggest that the oncogenic phenotype observed in cytokine-transformed cells is driven in part by H2B1O.

### H2B variants alter chromatin accessibility and up-regulate inflammatory and oncogenic pathways

We next asked if the observed changes in gene expression were associated with changes in chromatin accessibility and therefore performed ATAC-seq to identify differentially accessible regions (DARs). We found that DARs separated into distinct clusters by PCA, with cells expressing *HIST1H2BO* (and *HIST1H2BC*) grouping closer to cytokine-transformed cells (on principal component 1) than cells expressing *HIST1H2BH* (**Fig 7A**). Up-regulation of EMT factors and genes (from **Fig 6**) corresponded to increased accessibility at their corresponding gene promoters (**Fig 7B**). We thus annotated (DARs) at gene promoters, associated these annotations with proximal genes, and then compared genes associated with open or closed chromatin to up- or down-regulated genes, respectively. For all groups, there was significant overlap between open chromatin and up-regulated genes (30-48% of all DARs), and between closed chromatin and down-regulated genes (30-57% of all DARs; **Fig S9A**).

**Fig 7.**
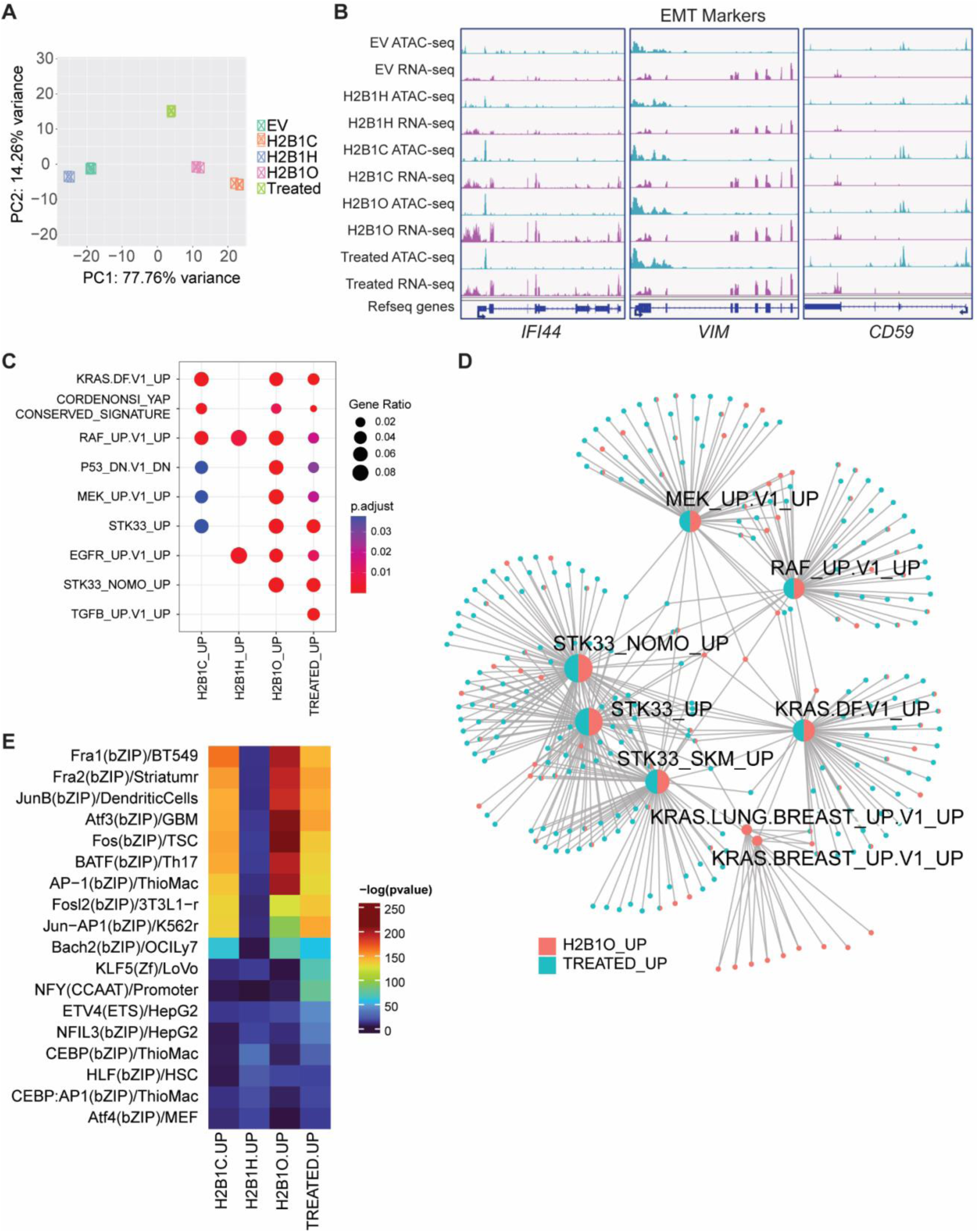
Cytokine-transformed MCF10A cells and cells expressing *HIST1B2BO* have overlapping ATAC-seq and RNA-seq signatures. A, Principal component analysis of ATAC-seq differentially accessible regions (DARs). **B**, Integrative Genomics Viewer (IGV) snapshots of ATAC-seq and RNA-seq peaks for genes involved in the epithelial-to-mesenchymal transition (EMT). cytokine-transformed cells and cells expressing *HIST1H2BC* and *HIST1H2BO* have similar, activated peaks/signatures. **C**, Over-representation analysis of activated MSigDB C6 oncogenic signatures that correspond with proximal changes in chromatin accessibility. Cells transfected with an empty vector (EV) served as the control. **D**, Enrichment map of the RAS-RAF-MEK pathway and overlapping genes in cytokine-transformed cells and cells expressing *HIST1H2BO*. Cells transfected with an empty vector (EV) served as the control. **E**, The top 18 DARs and up-regulated transcription factors in cytokine-transformed cells and cells transfected with H2B variant minigenes. Cytokine-transformed cells and cells expressing *HIST1H2BC* or *HIST1H2BO* exhibit increased chromatin accessibility and gene expression at the proto-oncogene AP-1 transcription factor family.

Next, we performed an over-representation analysis on the upregulated genes, correlating them with proximal increases in chromatin accessibility at the promoter regions (and vice versa). We found overlapping, enriched oncogenic and inflammatory pathways between cells expressing H2B variants and the cytokine-transformed cells (**Fig S9B**). Interestingly, the cytokine-transformed and *HIST1H2BO*-expressing cells shared eight proximally upregulated oncogenic signatures (**Fig 7C**); several MSigDB hallmark signatures (**Fig S9C**); and cell-substrate adhesion, cell-matrix adhesion, and embryonic body morphogenesis biological processes (**Fig S9D**) that are important in cancer cell migration and invasion [70]. We also noticed that cytokine-transformed and *HIST1H2BO*-expressing cells were enriched for many components of the RAS/RAF/MEK signaling cascade and its interactive STK33 pathway (**Fig 7D**), pathways that are associated with oncogenesis and tumor immune escape mechanisms [71, 72]. In contrast, cells expressing *HIST1H2BC* or *HIST1H2BH* were only enriched for a few components of this signaling pathway (**Fig S9E, F**). Furthermore, H2B1C, H2B1O, and the cytokine-transformed cells showed opened DARs for several AP-1 transcription factors that are associated with the most aggressive breast tumors, progressive inflammation, and poor clinical outcomes, including FosB, Fra-1, Fra-2, Jun-B, and BATF (**Fig 7E**; [73–77]). Collectively, these findings indicate that incorporating H2B1C, H2B1H, and H2B1O into chromatin induced differential chromatin accessibility changes, with H2B1O altering chromatin access and gene expression profiles involved in several tumorigenic pathways in ways that are comparable to cytokine (TGFβ/TNFα) transformed cells.

The relative overlap in oncogenic pathways and transcription factors between cytokine-transformed cells and cells expressing the H2B variants are consistent with the idea that the latter are undergoing EMT (see **Fig 5D, E**). We therefore asked if increasing H2B variant expression with cytokines (**Fig 4A** part **3**) would exacerbate or further drive MCF10A cells through the EMT. Cytokine treatment increased H2B variant expression, with the most notable increase seen in H2B1O (as expected; **Fig 8A**), and cytokine-treated cells expressing *HIST1H2BO* compared to cytokine-treated EV control cells were further enriched for pathways associated with the EMT, other oncogenic pathways, and signal transduction associated with tumor metastasis [78] (**Fig 8B** and **S9G)**. These data suggest that H2B1C, H2B1H, and H2B1O may epigenetically poise cells in different cellular states, and that further cytokine exposure might exacerbate cellular transformation.

**Fig 8.**
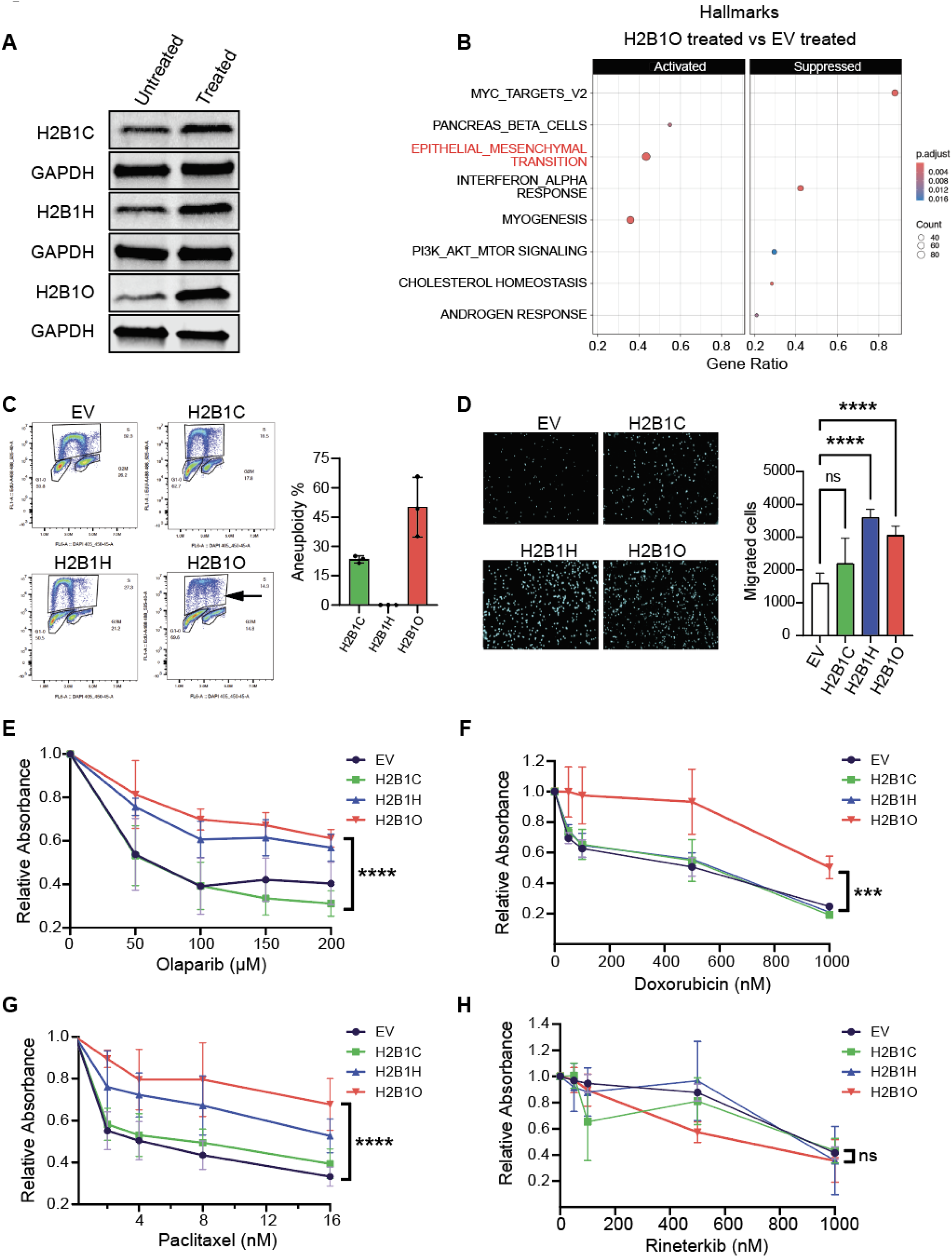
H2B1O makes cells resistant to common breast cancer chemotherapy agents. **A**, Immunoblot of H2B variant expression in untreated and cytokine-transformed cells. **B**, Enriched Hallmark signatures in cytokine-transformed cell expressing *HIST1H2BO.* **C**, Flow cytometry scatter plots and percent of transfected and cytokine-treated cells showing evidence of aneuploidy. The arrow indicates aneuploidy in the H2B1O flow cytometry data. Data in the bar plot are normalized to cells transfected with an empty vector and treated with TNFα and TGFβ. **D**, Number of migrating cells in a transwell assay (n = 3; biological replicates). Cytokine-treatment of transfected cells increases migration relative to cytokine-treated EV control cells (see **Fig 5D**). **E-H**, Response of transfected MCF10A cells to (**E**) Olaparib, (**F**) Doxoruicin, (**G**) Paciltaxel, and (**H**) Rineterkib, as measured with a colony formation assay.

### Incorporating H2BO into chromatin contributes to chemoresistance

During cell transformation, TGFβ cooperates with the oncogenic RAS and RAF signaling pathways to cause aneuploidy, which enhances tumor progression and chemotherapeutic drug resistance [79–85]. Incorporating H2B1C, H2B1H, and H2B1O into chromatin activates different components of the RAS/RAF/MEK pathway (**Fig 7D** and **S9E, F**), so we asked if these H2B variants can induce aneuploidy once cells are treated with TGFβ/TNFα. Stable expression of the H2B variants alone did not induce aneuploidy (see scatter plots in **Fig S5B**), but treating *HIST1H2BO*-expressing cells with TGFβ and TNFα did (**Fig 8C**). Stable H2B variant expression also increased cell migration relative to cytokine-induced transformation alone (**Fig 8D**). These data indicate that the cytokines cooperate with H2B variants to drive the EMT further, and that each H2B variant epigenetically regulates different inflammatory and/or oncogenic programs and differentially primes cells for the EMT.

We subsequently examined the responses of variant-transfected cells to commonly used chemotherapeutic drugs. Surprisingly, cells expressing *HIST1H2BO* were resistant to Olaparib (**Fig 8E**), Doxorubicin (**Fig 8F**), and Paclitaxel (**Fig 8G**), with cells expressing *HIST1H2BC* or *HIST1H2BH* showing variable resistance to each agent. Since incorporating H2B1C, H2B1H, and H2B1O into chromatin activates components of the RAS/RAF/MEK pathway (**Figs 7D**, **S9E,** and **F**), we then asked if Rineterkib (an RAF and MEK inhibitor), might be a more effective therapeutic agent. Indeed, each transfected cell type was sensitive to Rineterkib, especially those expressing *HIST1H2BO* (**Fig 8H**). These results suggest that *HIST1H2BO* over-expression might be a useful biomarker for guiding treatment decisions since breast cells over-expressing *HIST1H2BO* are resistant to standard chemotherapeutic agents but susceptible to a RAF/MEK inhibitor like Rinerterkib.

## Discussion

### Most H2B variants are dysregulated in breast cancer

H2B genes occur in three primary histone gene clusters (*HIST1*, *HIST2,* and *HIST3*), and *HIST1* gene expression is usually thought to occur only during cell division (i.e., replication-dependent histone production [86, 87]). However, some H2B variants in the *HIST1* cluster (including *HIST1H2BC*, *HIST1H2BD, HIST1H2BE,* and *HIST1H2BK*) are also expressed and spliced as polyadenylated mRNAs in terminally differentiated cells [88, 89], presumably as a mechanism to replace those histones that have a shorter half-life than their host cell. In fact, more recent studies are showing that many histone H2B variants across *HIST* gene clusters are replication-independent and can be polyadenylated [90, 91].

Prior studies have implicated some H2B variants in breast cancer, metastasis, and resistance to chemotherapy (reviewed in [92]). In this study, we found that 18 H2B variants are expressed in normal (presumably non-dividing) breast tissue and are dysregulated in their adjacent tumors. We were somewhat surprised that H2B1C (and its identical H2B1E, 1F, 1G, and 1I proteins) were not the predominant H2B variants in normal breast or tumor tissue (**Fig S1A**) given that so many genes encode for the same protein. We were also surprised that some H2B variants are preferentially over-expressed within breast cancer sub-types. *HIST1H2BO* over-expression, in particular, was associated with the most aggressive breast cancer sub-types and poor overall survival (**Fig 1**, and [93]). These data are consistent with the hypothesis that H2B variant expression in normal or cancerous tissue is a regulated, cell-specific process that is not always coupled to cell division [88], and that H2A-H2B dimers can be spontaneously exchanged with histones variants throughout the cell cycle [94, 95].

### H2B1O acts like an onco-histone

There are thousands of cancer-associated mutations in histone genes [96, 97], some of which occur at or near amino acids associated with post-translational modifications that regulate gene expression, and others that disrupt histone-DNA or histone-histone interactions (e.g., [98, 99]). For example, the H2B E76K mutation disrupts H2B-H4 interactions and forms unstable nucleosomes, especially at gene regulatory elements [100]. Nacev et al. (2024) also demonstrated that mutations in the arginine residues at positions 2 and 26 of the H3 N-terminus contribute to cancer by reducing the levels of H3K27me3, a histone modification associated with transcriptional repression [101]. These discoveries gave rise to the concept of an onco-histone, which is usually defined as a mutated histone that remodels chromatin to activate or suppress oncogenic pathways. An important realization from these (and related) studies is that seemingly minor histone amino acid substitutions can have profound consequences on nucleosome structure, stability, post-translational modifications, and gene expression. What is often overlooked, however, is that naturally occurring histone variants may also have unique biochemical, regulatory, and biological consequences.

In this study, we discovered that incorporating H2B1C, H2B1H, and H2B1O variants into chromatin cause breast epithelial cells to transition towards a mesenchymal phenotype. Indeed, most *HIST1H2BO*-expressing cells fully transitioned to the mesenchymal state and activated known oncogenic genes and pathways. H2B1O localized to promoters and activated EMT, BRCA, and KRAS pathways, while H2B1H deposition was enriched at the promoters of inflammatory response genes. Cells expressing *HIST1H2BC* and *HIST1H2BO* also had gene expression patterns that were more similar to cytokine-transformed breast epithelial cells than untreated cells (or those expressing *HIST1H2BH*). Thus, these H2B variants are enriched in different chromatin regions, associate with different oncogenic and inflammatory pathways, and are not functionally equivalent, even though they only differ from each other by one or two amino acids (at positions 2 and 39; see **Fig 3A** and [92]). Collectively, the data suggest that H2B1C and H2B1H epigenetically prime or poise epithelial cells for oncogenic transformation, whereas H2B1O acts like an onco-histone to drive epithelial cells through the EMT.

### H2B1O activates onco-inflammatory pathways that may create tumor vulnerabilities

H2B1O (in particular) activates many components of the RAS/RAF/MEK signaling pathway (**Fig 7D**), which may help explain why it is associated with the most aggressive breast cancers and low 5-year survival (**Fig 1C**). The RAS/RAF/MEK axis regulates cell proliferation, differentiation, migration, survival, and apoptosis (reviewed in [102]), and is often activated in breast cancer (especially EGFR+ and HER2+ breast cancers [103]) even when there are no RAS-related driver mutations [104]. Indeed, only ∼11% of all breast cancers have mutations in RAS/RAF/MEK/ERK pathway-related genes [105]. Treatment decisions often depend on what RAS/RAF/MEK/ERK mutations exist [106], but treatment response is difficult to predict and breast cancers often develop adaptive resistance to targeted therapies because of the complex feedback loops, pathway crosstalk, and post-translational modifications within the signaling network [107]. H2B1O activates the pathway in breast epithelial cells, leading to multi-drug resistance against standard breast cancer drugs ((**Fig 8E-H**); see also [71]). However, H2B1C, H2B1H, and H2B1O activate portions of the RAS/RAF/MEK/ERK pathway independent of any other driver mutations, such that transformed cells are susceptible to a RAF/MEK inhibitor (**Fig 8H**). The RAS/RAF/MEK/ERK cascade intersects with cell cycle, apoptosis, chemoresistance, immune-resistance, and immune-escape mechanisms; it is therefore possible that patients that do not have RAS-related mutations but over-express *HIST1H2BO* might benefit from a RAF/MEK inhibitor, either alone or in combination with other immuno- or chemotherapies. At the very least, *HIST1H2BO* expression should be considered in the same light as RAS-related DNA mutations [106] when devising treatment strategies.

### H2B1O is associated with the most aggressive cancers, race, and age

Clinical data consistently show that African American women exhibit a higher incidence of inflammatory breast cancer than white Americans [108, 109], and unequal response rates to chemotherapy even though they do not have an increased mutational burden [110–112]. These clinical data led to the hypothesis that there are biological differences in African American women that influence treatment response. In this study, we found that H2B1C, H2B1H, and H2B1O each “poise” or “prime” breast epithelial cells for the epithelial-to-mesenchymal transition (EMT), and that H2B1O can drive cells through the EMT on its own. *HIST1H2BO* expression is also highest in the most aggressive breast cancer sub-types; associated with decreased 5-year survival rates; enriched in young (< 45 years) African American and Asian patients; and up-regulates inflammatory and oncogenic pathways. It is therefore possible that H2B variant expression (rather than DNA mutations) might account for some of the racial disparities in breast cancer types and treatment response described in clinical studies. These studies also suggest that H2B variant expression might be used to sub-classify breast cancers and develop new therapeutic strategies for breast cancer patients who do not exhibit driver mutations in known oncogenes or histones.

## Methods

### Data collection from the TCGA database

TCGA gene expression data (based on GENCODE v29 gene annotations) were obtained using the recount3 v1.8.0 R package [113], with per-gene raw counts calculated using the ‘compute_read_counts’ function. Only samples with a library size >15 million reads were retained. Samples with the same sample ID were de-duplicated by retaining the sample with the highest number of uniquely mapping reads. Breast cancer molecular subtypes were obtained from [39] and refined using the metadata from the same publication; all samples with triple negative status were recoded to ‘TNBC’; basal tumor samples were labeled as ‘Basal_nonTNBC’; ‘LuminalA’ and ‘LuminalB’ samples were filtered for positive ER status by IHC; ‘HER2E’ samples were relabeled as ‘HER2+’ if both ER and PR status by IHC were negative; and samples not fulfilling any of the above criteria were removed unless they were labeled as “Metastatic” based on the recount3 metadata. Genes with <7 samples (number of samples of the smallest BRCA subtype) and >15 read counts were removed. Variance stabilizing transformed (VST) counts were obtained using the ‘vst’ function in DESeq2 v1.38.3 [114], with the parameter ‘blind=FALSE’ and the design set to ‘∼ Cancer_subtype’. The heatmap in **Fig 2A** was generated by plotting all expressed histone variant genes [115], histone chaperones [116], and genes related to chromatin modeling (GO:0006338). Sample clustering in the **Fig S2** heatmaps was based on the mean-centered variance stabilizing transformation counts for the genes shown in the heatmaps using K-means clustering with k set to 3.

### Patient survival **analysis**

The PrognoScan database provides various cancer microarray datasets that are combined with publicly accessible clinical annotations [40]. For this analysis, we used datasets for *HIST1H2BH* and *HIST1H2BO* expression and the R v4.4.0 package survival v3.6-4. Cox Proportional Hazard models were adjusted for histone expression level and run following the methods performed within PrognoScan. Variance stabilized transformed (VST) expression represents a continuous measure to estimate changes in hazard for a multiplicative change in expression level. K-means methods (stats v4.4.0 package) applied to VST expression generated categorical “low” vs. “high” expression levels for survival models, thus estimating the change in hazard between patients with low and high expression levels. The emmeans (v1.10.1) package was then used to extract hazard ratio “low” vs. “high” contrasts. Cohort-specific information is accessible for overall survival within a cancer type through the PROBE ID section in the PrognoScan query return value.

### Cell Lines

MCF10A human breast epithelial cells were obtained from the American Type Culture Collection (ATCC). This cell line was cultured as previously described [117] in phenol red-free DMEM/F12 media supplemented with 5% horse serum (Gibco), 1X penicillin-streptomycin (Gibco), 20 ng/mL EGF (Peprotech), 0.5 μg/mL hydrocortisone (Sigma-Aldrich), 100 ng/mL cholera toxin (Sigma-Aldrich), and 10 μg/mL insulin (Sigma-Aldrich). The human breast cancer cell lines MCF7 (metastatic luminal A), T47D (luminal A), BT474 (luminal B), and MDA-MB-231 (TNBC) were obtained from ATCC. MDA-MB-231 was cultured in phenol red-free, high glucose DMEM (Gibco) supplemented with 10% FBS (Gibco). T47D cells were cultured in RPMI 1640 medium (Gibco) supplemented with 10% FBS (Gibco). MCF7 and BT474 were cultured in MEM medium (Gibco) supplemented with 10% FBS (Gibco), 1% MEM Amino Acids Solution (Gibco), 1% MEM Non-Essential Amino Acids Solution (Gibco), 1% Sodium pyruvate (Gibco), and 10 μg/mL insulin (Sigma-Aldrich). All cell lines were maintained at 37°C and 5% CO_2_ and were tested for mycoplasma contamination using the Mycplasma PCR Detection Kit detection kit (Invitrogen). To induce EMT with TGFβ and TNFα, cells were treated with 5 ng/mL of recombinant human TGF-β1 (PeproTech) and 5 ng/mL of recombinant human TNFα (PeproTech) for 6 days. During treatments, cells were split every other day.

### Minigene systems and stable expression cell lines

We designed minigene systems as described in Saintilnord et al. (2024), with each minigene containing 2xHA-tagged H2B variants that were controlled by their endogenous promoters [38]. These minigene plasmids and an empty vector (EV) control were custom synthesized by GeneScript (Piscataway, NJ) and inserted into a puromycin selection promoter-less pcDNA3.1 vector at the NDeI/ BamHI restriction sites. Stable cell lines containing an empty vector or HA-labeled H2B variant were derived by electroporating ∼1 x 10^6^ MCF10A cells and 2 µg of the minigene plasmid with a Lonza 4D-Nucleofector^®^ X Unit (Lonza) and SF Cell Line 4D-Nucleofector™ X Kit S (Lonza) according to the manufacturer’s instructions. Cells were then transferred from nucleofection strips into 6-well plates with pre-warmed growth media and grown for 48 h at 37°C and 5% CO_2_. Thereafter, we added 1.5 μg/mL puromycin (Gibco^TM^) to the growth media and cultivated the cells for 16 weeks before using them for subsequent experiments.

### Histone octamer preparation

Histone octamers were obtained from the Histone Source at Colorado State University, where they were purified according to established protocols [118, 119]. The lyophilized histones were resuspended at 3-5 mg/mL for 1 h in an unfolding buffer (20 mM Tris-HCl pH 7.5, 7M guanidinium, and 10 mM dithiothreitol). Suspensions were then centrifuged to remove insoluble aggregates, and the concentration of each unfolded histone was determined by measuring absorbance at 276 nm.

Heterodimers of H2A and H2B were refolded at a 1:1 ratio, as were tetramers of H3 and H4. Histone octamers were subsequently formed as previously described [38] by combining the re-folded heterodimers (H2A and H2B) and tetramers (H3 and H4) in a 1.2:1 ratio, and performing double dialysis into a refolding buffer (10 mM Tris-HCl pH 7.5, 1 mM EDTA, 2M NaCl, and 5 mM β-mercaptoethanol). Octamers were then purified on a Superdex 200 gel filtration column (GE Healthcare), and the purity of each octamer confirmed by SDS-PAGE in 18% acrylamide gels (29:1 acrylamide: bisacrylamide).

### Nucleosome positioning sequence DNA preparation

A plasmid containing the Widom 601 nucleosome positioning sequence was constructed following previously described protocols [38] by adding 50 bp of linker DNA at each end of the NPS. This 247 bp DNA construct was labeled with a radioactive isotope α-^32^P (Revvity, Waltham, MA) via PCR amplification. The amplified DNA was concentrated by centrifugation with a 30K Amicon® Ultra-15 Centrifugal Filter Unit (Millipore Sigma, Burlington, MA). The concentrated DNA was then run on a 14% native polyacrylamide gel, and the 247 bp band excised, crushed, and incubated on a rotator in Crush-N-Soak buffer (0.75M ammonium acetate, 0.3M sodium acetate, 1 mM EDTA pH 8.0, 0.1% SDS) at room temperature overnight. DNA was further extracted by butanol reduction and buffer exchange into 0.5X TE (5 mM Tris, 0.5 mM EDTA pH 8.0). Labeled, purified DNA was again concentrated with a 30K Amicon® Ultra-4 Centrifugal Filter Unit and then dialyzed overnight at 4°C into 0.5X TE using a 10 kDa Slide-A-Lyzer Mini Dialysis Unit (# 0069570, ThermoFisher Scientific). The concentration of purified DNA was then determined using a Thermo Scientific™ NanoDrop™ OneC Microvolume UV-Vis Spectrophotometer (Thermo Scientific™ 840274200).

### Preparation and reconstitution of mono-nucleosomes

Nucleosomes were prepared and purified according to previously established protocols [38, 119, 120]. Briefly, ^32^P-labeled NPS DNA was mixed with purified and refolded histone octamers at a ratio of 1.25:1, and nucleosomes reconstituted by double dialysis. The first dialysis was into a high-salt buffer (2M NaCl in 0.5X TE, 1 mM benzamidine hydrochloride) at 4°C for 5-6 h, and then into a low-salt buffer (0.5X TE, 1 mM benzamidine hydrochloride) at 4°C for another 5-6 h. DNA was then dialyzed overnight at 4°C against 4L of fresh low-salt buffer. Re-constituted nucleosomes were purified on a 5-30% w/v sucrose gradient that was prepared from pre-chilled sucrose solutions using a BioComp Gradient Station, as per the manufacturer’s instructions. Nucleosome sucrose gradients were then centrifuged at 41,000 rpm and 4°C for 22 h in a SW41 Ti rotor in an Optima L-100 XP ultracentrifuge (Beckman Coulter, Brea, CA). After centrifugation, sucrose gradients were fractionated into 0.4 mL aliquots and analyzed by 5% native polyacrylamide gel electrophoresis. Fractions that contained centrally positioned nucleosomes were concentrated into 0.5X TE at 4°C using a 30K Amicon® Ultra-15 Centrifugal Filter Unit (#UFC903024, Millipore Sigma, Burlington, MA). To confirm the purity of the isolated nucleosomes, they were analyzed on native 5% acrylamide gels. The gels were stored for 24 h on a phosphor imaging screening and imaged the following morning using a Typhoon biomolecular imager (Cytiva). Purified nucleosomes were also precipitated with 5 mM MgCl_2_ [121], heated at 90°C in 1× SDS loading buffer, separated by SDS-PAGE on an 18% gel, and stained with Coomassie blue. Band intensities were then measured using a ChemiDoc MP Imaging System.

### Mono-nucleosome enzyme accessibility assays

Mono-nucleosomes containing ^32^P-labelled NPS DNA were digested with either MNase, DNAse I, or Exonuclease III. For micrococcal nuclease (MNase), ∼200 ng of mono-nucleosomes was incubated with 0.003125 U/µL of MNase. At the designated time intervals, 100 µL aliquots were removed into a new tube containing 20 µL of stop solution (0.5 M EDTA, 0.2 M EGTA, 10% SDS, and 0.25 µg/µL of proteinase K), as previously described [38]. The digested DNA was then purified with a QIAquick PCR Purification Kit (Qiagen, Hilden, Germany), eluted in 40 µL of 0.5X TE, mixed with Orange G (Sigma Aldrich), and analyzed by native PAGE on 5% acrylamide gels. For DNAse I and Exonuclease III assays, 50 ng of ^32^P-labeled mono-nucleosomes were treated with either 0.00625 U/µL of DNase 1 or 2 U/µL of Exonuclease III. At the designated time intervals, 15 µL aliquots were transferred to new tubes containing 2 µL of stop solution (20 mM EDTA, 0.2% SDS, and 0.25 µg/µL of proteinase K). An equal volume of 2X formamide loading buffer was then added to each tube, and the contents incubated for 15 min in a boiling water bath. The digested DNA was separated by denaturing PAGE (8M urea) on 10% acrylamide gels. Gels were then wrapped in plastic, placed on a phosphor screen overnight, imaged using an Amersham Typhoon (GE Healthcare), and quantified with ImageQuant software.

### DNA preparation for nucleosome arrays

A plasmid containing a 17-mer repeat of the 601 NPS was a kind gift from the Poirier lab (see [122]). The plasmid was transformed into DH5α competent cells and purified with a ZymoPure II Plasmid Maxiprep Kit. Purified plasmid was digested with DdeI (NEB, Ipswich, MA) overnight at 37°C, resulting in an ∼3 kb DNA fragment. Digested plasmid DNA was then purified using phenol-chloroform extraction and analyzed on a 0.7% agarose gel to confirm purity and digestion efficacy.

### Reconstitution of nucleosome arrays

Nucleosome arrays were reconstituted as described above for mono-nucleosomes, except that histone octamers and the 17-mer NPS DNA were combined in a 1:0.9 ratio. Purified nucleosome arrays were subsequently isolated on 5-40% w/v sucrose gradients in 0.5X TE by centrifuging the gradients at 39,000 rpm and 4°C for 8 h in a SW41 Ti rotor (Beckman Coulter, Brea, CA). After centrifugation, sucrose gradients were fractionated into 0.4 mL aliquots (as above) and analyzed on 1% agarose gels. Fractions containing the nucleosome arrays were concentrated on a 30K Amicon filter unit into 0.5X TE. The concentration of nucleosome arrays was determined based on DNA content, as measured with a NanoDrop spectrophotometer. Nucleosome array quality was evaluated by precipitating ∼100 ng of each array with 5mM MgCl_2_, heating to 90°C in 1X SDS loading buffer, performing SDS-PAGE, and staining with Coomassie blue. Arrays were also digested into monomers with AvaI (NEB) and analyzed by 1% agarose gel electrophoresis.

### Restrictions enzyme assays for nucleosome arrays

Nucleosome arrays (∼150 ng) were digested with either 50 units of BamHI (NEB) for 1 h or 10 units of PstI (NEB) for the indicated time points. The digestion reactions were stopped with 1.5 mM EDTA (pH 8.0), digested with proteinase K (1µg/µL) for 15 min at 37°C, and analyzed on 0.7% agarose gels.

### Atomic Force Microscopy (AFM)

Nucleosome arrays were prepared as described in Saintilnord et al. (2024) by first mixing the 17-mer NPS DNA fragment with histone octamers at a 1:0.9 ratio and incubating on ice for 30 min in (2M NaCl, 10 mM Tris HCl pH 8, 1 mM EDTA) [38]. The mixtures were then transferred to 3.5 kDa Slide-A-Lyzer Mini Dialysis Units (#0069558, ThermoFisher Scientific) and dialyzed at 4°C against 1X TE containing various NaCl concentrations: 1M NaCl for 2 h; 0.8M NaCl for 2 h; 0.6M NaCl overnight; and 0.15M NaCl for 2 h. The purified nucleosome array was diluted 40-fold into (35 mM KCl, 15 mM NaCl, 5 mM HEPES, and 0.25% glycerol), and then deposited in a single layer on freshly cleaved mica that had been functionalized with aminopropyl-silanetrane (APS) [38, 123]. AFM images were acquired on an Oxford Instruments Asylum Research’s Cypher S AFM equipped with silicon cantilevers (Olympus OTESPA-R3 cantilevers; nominal resonances of ∼300 kHz and a stiffness of ∼42 N/m), operating in non-contact tapping mode in ambient air. Image processing was performed as previously described [123], with images analyzed in ImageJ. Statistical analyses and data visualization were performed in GraphPad Prism version 10 (La Jolla, CA, USA) [38].

### Flow cytometry

MCF10A cells (∼2 x 10^6^) that were transfected with an empty vector or H2B minigene systems were seeded into fresh culture medium for 24 h. We used the Click-it^TM^ Plus EdU Alexa Fluor^TM^ 488 Flow Cytometry Assay Kit (Invitrogen) to detect the expression of H2B variants across all cell cycle phases, following the manufacturer’s instructions with the modifications detailed below. Cells in S-phase were stained by adding 10 μM of EdU to (from the Click-it Plus kit) to the culture medium and cells were incubated for 2 h at 37°C and 5% CO_2_. Cells were then harvested using trypsin, washed with 1% BSA in phosphate buffered saline (PBS), and counted to ensure the same number of cells were used for each subsequent staining condition. Cells were then fixed with 70% ethanol for one hour on ice, pelleted, and washed with 1% BSA in PBS. Cell pellets were dislodged and permeabilized with 100 μL of 1X Click-iT^TM^ permeabilization and wash reagent for 15 minutes. For EdU detection, 0.5 mL of Click-iT^TM^ Plus reaction cocktail was mixed into each tube of fixed cells and incubated for 30 min. Cells were then washed with 1X Click-iT^TM^ permeabilization and wash reagent. To stain for intracellular antigens, cell pellets were resuspended in 100 μL of 0.1% Tween 20 in PBS (PBST). The cells were then incubated with 3 μg/mL of HA-Tag primary antibody (C29F4, Cell Signaling) and 3 μg/mL of conjugated H3 pSer10 antibody (Alexa Fluor® 594, BioLegend), which served as a marker for M phase, for a duration of 30 min. Cells were then washed with PBS, and incubated for 1h in 100 μL PBST with secondary anti-rabbit IgG (H+L) (Alexa Fluor^®^ 647 Conjugate, Cell Signaling). Stained cells were then washed twice with PBST and resuspended in 300 μL of PBS with 5 μg/mL DAPI to stain cells for DNA content.

After staining, cell suspensions were transferred to FACS tubes and analyzed on a BD Accuri B6 flow cytometer (BD Biosciences). The gating strategy was designed to identify cells in each phase of the cell cycle. G0/G1- and G2 phase cells were distinguished by DAPI staining; S-phase cells were identified by EdU staining; and M cells were identified by H3 pSer 10 staining. Data were analyzed in the FlowJo software package.

### RT-qPCR

Total RNA was isolated using the Quick-RNA Miniprep Kit (Zymo Research) according to the manufacturer instructions, except that contaminating DNA was first digested with (RNase-free) DNase I. Purified RNA was quantified by measuring the absorbance at 260 nm using a NanoDrop spectrophotometer (Thermo Fisher Scientific). cDNA was synthesized using the iScript cDNA synthesis kit (Bio-Rad), and quantitative expression of target genes were analyzed by real-time quantitative PCR (RT-qPCR) using SYBR green master mix (Life Technologies) on a Real-Time PCR system (Bio-Rad Thermal Cycler). Target gene expression was normalized to *GAPDH* and TATA-binding protein (*TBP*) expression. Primer sequences are listed in materials and resources table.

### Whole cell protein extraction and immunoblot

Approximately 3 x10^6^ cells were thawed and resuspended in 200 μL 1X RIPA lysis buffer (Thermo Fisher Scientific) with Halt™ Protease Inhibitor Cocktail (Thermo Fisher Scientific). Cells were then sonicated for 12 cycles with 30 sec ON and 30 sec OFF in Bioruptor*Plus* sonicator, then centrifuged for 15 min at 13,000 RPM at 4°C. The supernatant was transferred to a clean Eppendorf tube, and protein concentration measured with a Pierce BCA Protein Assay Kit (Thermo Fisher Scientific) and EnVision plate reader according to the manufacturer’s instructions. For immunoblots, ∼50 μg total protein from each sample was boiled for 10 min in 1X SDS PAGE protein loading buffer and then separated by electrophoresis through 4-20% gradient gels (Bio-Rad). The separated proteins were transferred to a nitrocellulose membrane (iBlot Transfer Stack, Invitrogen) using an iBlot^TM^ 2 Gel Transfer Device (Invitrogen). Blots were then blocked in 5% fat free dry milk, and proteins of interest detected with the target antibodies.

### Subcellular protein fractionation

Approximately 3 x10^6^ cells were processed through a Subcellular Protein Fractionation Kit for Cultured Cells (Thermo Fisher Scientific) as per the manufacturer’s instructions. Protein fractions were subsequently analyzed by immunoblot, as described above.

### Transwell migration assay

MCF10A cells transfected with HA-tagged H2B variants or empty vector (EV) were seeded in cell culture flasks to reach ∼50% confluency. Culture media was then aspirated, cells were washed with PBS, starved overnight by adding blocking/starving solution (DMEM/F12 without supplements), and placed back in the incubator at 37°C and 5% CO_2_. The next day, 24-well transwell plates and 8 μm pore size Boyden chamber inserts (BD Biosciences) were pre-coated with collagen solution and placed in the incubator for 1 h. Wells and inserts were then washed with PBS and blocked with blocking/starving media for at least 1 h. Starved cells were recovered by trypsin digestion, quenched with blocking/starvation solution, pelleted by centrifugation, resuspended in blocking/starvation solution, and counted with an automated cell counter (Invitrogen). Blocking/starving media was then aspirated from wells and inserts, and migration-inducing media (growth DMEM/F12 media) was added to the lower chamber. Approximately 5 x 10^4^ cells (in 100 μL blocking/starving suspension) were added to the upper chamber of the inserts and incubated for 24 h. The incubation time was reduced to 10 h for those cells that were previously treated (and transformed) with TGFβ and TNFα. After incubation, non-migrating cells on top of the insert were wiped off with wet cotton swabs, and migrated cells remaining on the lower surface of the membrane were fixed with 100% methanol for 45 min. Inserts were then dipped in water, aspirated to remove excess water, stained with (10 μg/mL DAPI and 1% TritonX-100 in PBS) for 30 min, then rinsed in PBS. Images of stained cells were captured with an Evident Olympus microscope at 40X magnification. Quantification was carried out in an automated way using the cellSens counting module. Images of the area covered with DAPI stained nuclei were generated using a threshold and settings that were appropriate for control unstained samples, and these settings were used throughout the analysis.

### Crystal violet assay for chemotherapeutic drug response

Cells were seeded into 96-well plates and treated the next day with fresh media containing 0.1% DMSO (vehicle control), Olaparib (0-200 μM), Paclitaxel (0-16 nM), Doxorubicin (0-1000 nM), or Rineterkib (0-1000 nM). Chemotherapy media was replaced every other day for 4 days, after which cells were fixed in 4% formaldehyde (Electron Microscopy Sciences; in 1X PBS) for 30 min. The fixative was removed, plates were washed with PBS, and cells stained with 1% crystal violet 15 min. The staining solution was then removed, and the plates washed three times under running water. Plates were allowed to dry at room temperature, the residual crystal violet stain eluted with 100 μL of 100% methanol, and absorbance measured at 562 nm with an EnVision plate reader (Perkin Elmer). Each experiment had four biological and two technical replicates.

### Total RNA-seq

Total RNA was isolated as described above, and RNA quality assessed with a BioAnalyzer. Library preparation and total RNA-seq were conducted by the Van Andel Research Institute Genomics Core. After depleting ribosomal RNAs with the QIAseq FastSelect –rRNA HMR Kit (Qiagen, Germantown, MD, USA) from 500 ng of total RNA, RNA was fragmented to 300-400 nt and libraries generated using the KAPA RNA HyperPrepKit (Kapa Biosystems, Wilmington, MA USA) as per the manufacturer’s protocol. cDNA fragments were then ligated to IDT for Illumina TruSeq UD Indexed adapters (Illumina Inc, San Diego CA, USA), and the libraries amplified by PCR. The quality and quantity of the finished libraries were assessed using Agilent DNA High Sensitivity chips (Agilent Technologies, Inc.), a QuantiFluor® dsDNA System (Promega Corp.,Madison, WI, USA), and Kapa Illumina Library Quantification qPCR assays (Kapa Biosystems). Individually indexed libraries were then pooled and sequenced (50 bp paired-end) on an Illumina NovaSeq6000 sequencer to an average depth of 50M raw paired-reads per transcriptome.

Base calling was done by Illumina RTA3, and the output of NCS was demultiplexed and converted to FastQ format with Illumina Bcl2fastq v2.20.0. We used TrimGalore v0.6.10 and parameters ‘--paired -q 20’ to trim reads and remove low-quality bases and adapter sequences. Trimmed reads were then aligned to the hg38 reference genome that was downloaded from GENCODE release 33 [124]. Gene counts were obtained using STAR v2.7.10a with the parameters ‘--twopassMode Basic --quantMode GeneCounts’ [125]. Raw counts were converted to variance stabilizing transformation (VST) counts. Principal component analysis (PCA) was done on the top 10,000 genes with the highest variance using variance as calculated by the vst() function in DESeq2 v1.40.2 with ‘blind=FALSE’ [114]. We used DESeq2 and the design ‘∼ group’ for differential expression analysis. Pairwise contrasts were deemed significant at an adjusted p-value ≤ 0.01. Log fold changes were shrunk using the DESeq2 lfcShrink function with the parameters, ‘type=“ashr”’. GSEA was conducted using clusterProfiler v4.6.2 with the parameter ‘eps = 0.0’ [126], ranking genes by the shrunken log-fold changes and significance cutoff of adjusted p-value ≤ 0.05. MSigDB annotations were retrieved using msigdbr v7.5.1, and summary plots were created using enrichPlot v1.20.3.

### ATAC-seq

MCF10A cells transfected with empty vector, *H2B1C*, *H2B1H*, or *H2B1O* minigenes were pelleted, washed, counted, and frozen in growth media containing 45% horse serum and 5% DMSO. Cryopreserved cells were then sent to Active Motif (Carlsbad, CA) for ATAC sequencing as previously described [127], with some modifications [128]. Briefly, cells were thawed, pelleted, and washed with cold PBS. Then, ∼1×10^5^ cells were resuspended in lysis buffer, and the nuclei pelleted by low-speed centrifugation. Genomic DNA was then tagmented using a Nextera Library Prep Kit (Illumina), and tagmented DNA fragments purified through a MinElute PCR purification kit (Qiagen). Purified DNA was amplified with 10 PCR cycles, and the amplified DNA purified using Agencourt AMPure SPRI beads (Beckman Coulter) as per the manufacturer’s instructions. Resulting libraries were quantified using the KAPA Library Quantification Kit specifically for Illumina platforms (KAPA Biosystems) and subjected to 42-bp paired-end sequencing on a NextSeq500 sequencer (Illumina). Illumina base-calling and de-multiplexing was done using bcl2fastq2 conversion software (v2.20). Reads (in FASTQ format) were processed through skewer v.0.2.2 to eliminate adaptor sequences, and Samtools (v0.1.19) was used to remove duplicates. Reads were then aligned to the human genome (hg38) using the BWA algorithm (mem mode; default settings) [129]. The sequencing reads were treated as matched pairs, and only uniquely mapped reads with a mapping quality score ≥1 were retained. Alignments were extended *in silico* at their 3’-ends to a length of 200 bp and assigned to 32-nt bins along the genome. The genomic “signal maps” were then converted to bigwig files using deeptools (v3.5.1).

For comparative analysis, we normalized data by reducing the number of usable tags for each sample in a group to match the lowest number of usable tags among all samples in that group. We used the MACS 2.1.0 algorithm [130] with of a p-value cutoff of 1e-7, no control file, and with the ’--nomodel’ option for peak calling. Any peaks that were on the Encyclopedia of DNA Elements (ENCODE) blacklist [131] of known false peaks were removed. Signal maps and peak locations were used as input data for Active Motif software, which generates Excel tables that compare peak metrics between samples, peak locations, and gene annotations. Signal tracks were visualized using Integrative Genomics Viewer (IGV) [132].

The outputs from the Active Motif software were used to annotate peaks to the nearest genomic features using the annotatePeak function in ChIPseeker v1.34.1 with the parameters ‘tssRegion = c(-3000, 500)’ [133]. The peaks were also annotated to ENCODE3 predicted cis-regulatory elements (cCRE) using the GenomicRanges v1.50.2 software and a minimum overlap of 1 base pair. Differentially accessible peaks were chosen based on an adjusted p-value < 0.1.

### ATAC-seq and RNA-seq integration

Significantly up- or down-regulated genes were compared to genes with significantly opened or closed promoters (annotated as described above), respectively. The lists of differentially expressed genes with matching differentially accessible promoters were used for over-representation analysis using the compareCluster function in clusterProfiler v4.6.2 with a default adjusted p-value ≤ 0.05. The package msigdbr was used to retrieve MSigDB annotations (as described for the RNA-seq analysis), and the summary plots of the enrichment results were created using enrichPlot v1.18.4.

### CUT&RUN

Approximately 5 x 10^5^ cells per reaction were used to prepare CUT&RUN libraries with a CUTANA kit, with minor modifications. Briefly, cells were pelleted by low-speed centrifugation at 4°C, washed twice with 100 μL of Wash buffer, and then resuspended in 105 μL Wash buffer. Separately, ConA beads were activated by transferring 11 μL of ConA beads into an 8-strip tube, washed twice with 100 μL of cold Bead Activation Buffer as per the manufacturer’s instructions, and then resuspended in 11 μL of the same buffer. A volume of 100 μL of the cell suspension was then mixed with the activated ConA beads in the 8-strip tube. The bead/cell slurry was incubated for 10 minutes at room temperature on a nutator to adsorb cells to the beads and then placed on ice. After the slurry cleared, the supernatant was removed, and the beads/cells were resuspended in 50 μL of cold Antibody Buffer.

For reactions targeting H3K4me3 positive and IgG negative control antibodies, 2 μL of K-MetStat Panel and 1 μL of antibody per reaction were added. For other target antibodies, 1 μL of HA-tag (Cell Signaling, C29F4) and 1 μL of H3K27me3 (Epicypher, SKU: 13-0055) antibodies were added per reaction. All antibodies were thoroughly mixed with the bead/cell slurry via pipetting and incubated overnight at 4°C. The following day, the 8-strip tube was placed on a magnetic stand, and the supernatant was removed. Subsequently, cells were washed in 200 μL of cold Cell Permeabilization Buffer (Wash buffer + 0.01% Triton X-100), resuspended in 50 μL of the same buffer, gently vortexed, and placed on ice.

For enzymatic digestion, 0.4 μL of pA-MNase was added per reaction, and the mixture was incubated for 10 minutes at room temperature on a nutator. The supernatant was subsequently removed using magnetic beads, the samples were washed four times with Cell Permeabilization Buffer and then resuspended in 50 μL of the same buffer. To facilitate chromatin digestion and release, 1 μL of 100 mM calcium chloride was added to each reaction and the mixture incubated on the nutator for 2 h at 4°C. Subsequently, 33 μL of Stop Buffer and 1 μL of *E. coli* spike-in DNA were added per reaction, and the samples placed in a thermocycler at 37°C for 10 min.

Using magnetic beads, the slurry was collected and transferred to 1.5 mL tubes for DNA purification. To each reaction, 420 μL of DNA Binding Buffer was added and mixed by vortexing. Each reaction was then processed through a DNA Cleanup Column by centrifuging at 16,000 x g for 30 sec at room temperature. The flow-through was discarded, and the columns were washed twice with 200 μL of DNA Wash Buffer. An additional spin was performed to ensure the column was completely dry before transferring it to a clean 1.5 mL tube. Finally, 12 μL of DNA Elution Buffer was added to the center of the column, incubated for 5 min, and then centrifuged at 16,000 x g for 1 min at room temperature. DNA quantity and quality were evaluated using an Agilent TapeStation prior to library prep.

Library preparation and sequencing were conducted by the Fred Hutchinson Cancer Center Genomics Core. The end repair, A-tailing, adapter ligation, post-ligation cleanup, library amplification, and post-amplification cleanup steps were all performed using a KAPA HyperPrepKit (Kapa Biosystems, Wilmington, MA USA) as per the manufacturer’s protocol. The quality and quantity of the finished libraries were again evaluated using a TapeStation. We used “Region View” in the TapeStation analysis software to select the library’s upper and lower boundaries, and to avoid adapter-dimer peaks (∼150 bp). Individually indexed libraries were then pooled and sequenced on an Illumina NovaSeq6000 sequencer.

FastQC [134] was used to examine read quality, and high-quality reads were then aligned to the reference genome (hg38) using Bowtie2 [135]. Genome coverage profiles were generated using bedtools genomecov [136], and MACS2 [130] was used to call peaks. MACS2 was run in both target-only and IgG-controlled (using all IgG reads) modes. For H3K4me3 narrow peaks, IgG-controlled peaks were kept for downstream analysis if they overlapped with treatment-only peaks, the q-value was < 0.01, and the enrichment was > 2. For broad H3K27me3 peaks, target-only peaks were kept for downstream analysis if they overlapped with IgG-controlled peaks, the q-value was < 0.01, and the enrichment was > 2. For HA-tagged H2B proteins, MACS2 broad peaks with IgG control and EV-HA as a negative control were kept if they overlapped with target-only peaks, q-value < 0.01, and the enrichment was > 2. Peaks present in only one replicate of a condition were excluded, and peaks from all conditions were pooled and merged into a set of non-overlapping consensus peaks. FeatureCounts [137] in Subread 2.0.0 was then used to count reads mapped to each consensus peak.

Differential peaks between samples were detected using edgeR 3.36.0 [138] and the filterByExpr function with min.count = 10 and min.total.count = 15. We used the TMM method [97] to normalize the resulting count matrix, and the quasi-likelihood method to test for significance. Significantly differential peaks were defined as those with >1.5-fold change (in either direction), and a Benjamini Hochberg-corrected FDR < 0.05. BEDTools v2.30.0 [136] was used to identify genes where H3K4me3 or H2B variants occurred within -1 Kb to +100 bp of a TSS, and where H3K27me3 occurred within 2.5 Kb of TSS. Pathway enrichments were identified as described above for the ATAC-seq and RNA-seq integration data but using clusterProfiler v4.13.0 and enrichPlot v1.25.0.

## Data availability

The accession number for the raw RNA sequencing data reported in this paper is GEO: XXXX, for the raw CUT&RUN sequencing data reported in this paper is GEO: XXXX and for the raw ATAC sequencing data reported in this paper is GEO: XXXX.

## Acknowledgments

We thank members of the Fondufe-Mittendorf for critical feedback on the experimental designs. We thank Van Andel Research Institute Genomics, Flow Cytometry, Bioinformatics and Biostatistics cores for valuable technical assistance. We thank the Histone Source at Colorado State University for assistance with histone purification. We thank Feinan Wu and the Fred Hutchinson Cancer Center Genomic & Bioinformatics core for assistance with CUT&RUN sequencing, which is supported by the Bioinformatics Shared Resource of the Fred Hutch/University of Washington/Seattle Children’s Cancer Consortium (P30 CA015704). This study was supported by the National Science Foundation grant NSF/MCB 016515 (YFM), National Institutes of Health grants R01ES024478 (YFM), R01ES034253 (YFM), and 1R01ES036051-01 (YFM), the Van Andel Institute institutional support (YFM), and by NCI grant CA266078 (JDL).

## Author contributions

H.D generated H2B variants expressing cells and performed experiments utilizing them, as well as RNA-seq, CUT&RUN-seq, ATAC-seq, and biophysical experiments. K.H.L performed TCGA transcriptomics and the other sequencing analyses. W.N.S performed AFM experiments with the assistance of D.P.M. H.N.D performed survival analysis and population distribution among patients. E.L assisted with sequencing analysis of chemically treated samples. F.R.P performed RT-qPCR of histone variants expression across different cell lines. H.D and Y.F-M designed experiments and conceptualized the project with the assistance of M.G.B, J.D.L, and Y.D. H.D, D.P.C and Y.F-M assisted with data interpretation, and manuscript and figure editing. H.D. wrote the manuscript with input from D.P.C and Y.F-M. All authors have reviewed and provided editing suggestions for the manuscript. All diagrams were designed with Biorender.com.

## Materials and Resources

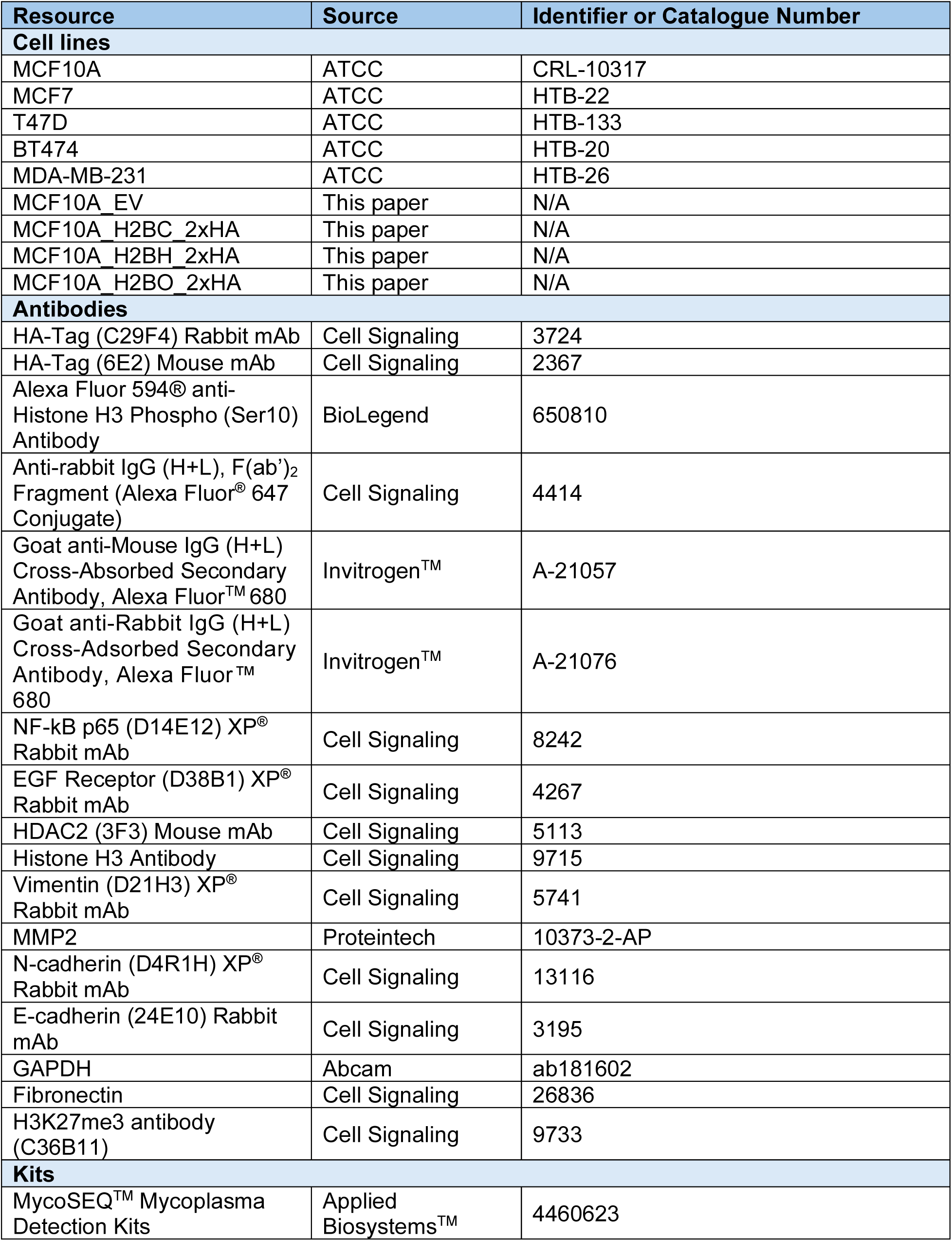
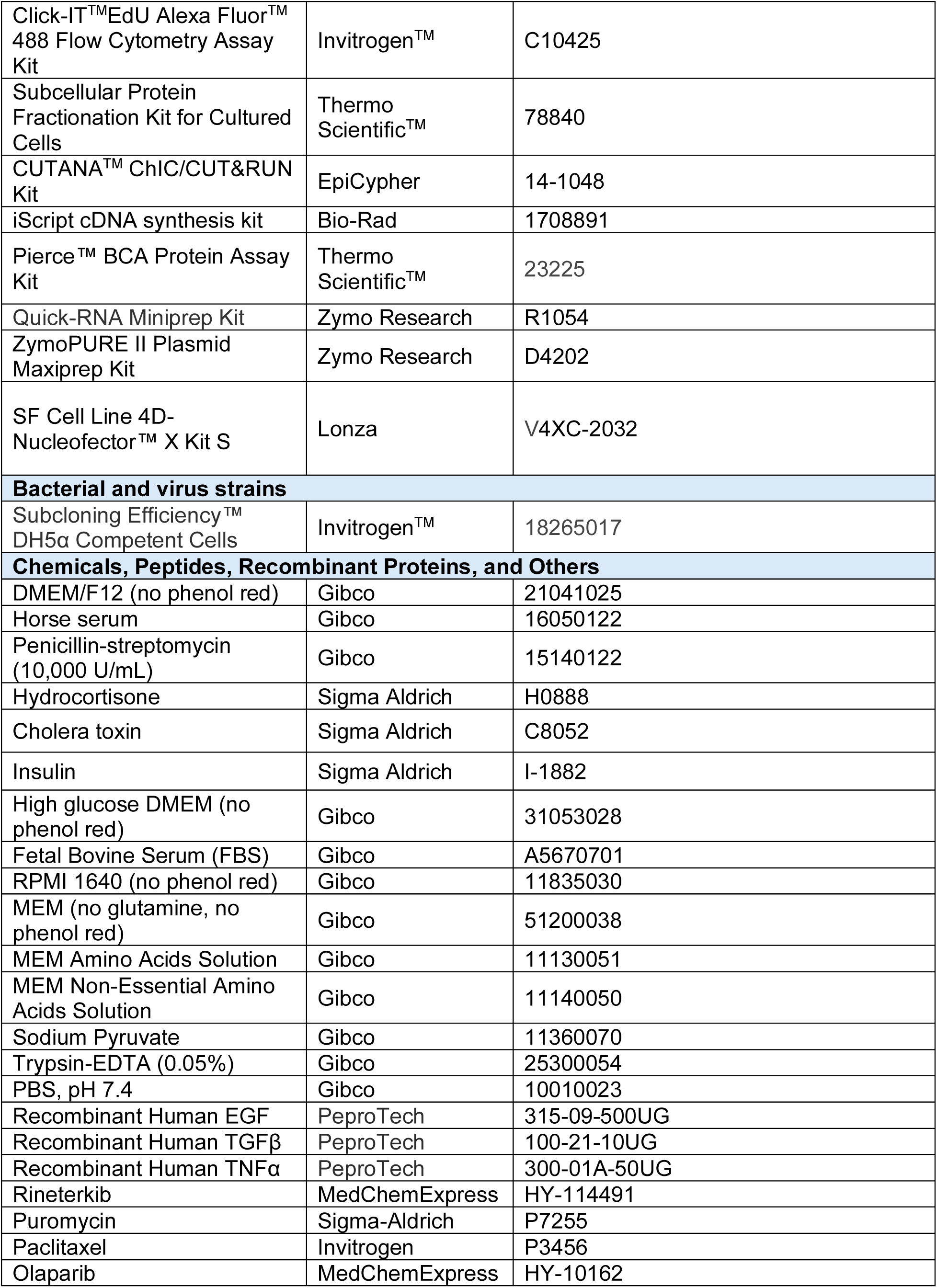
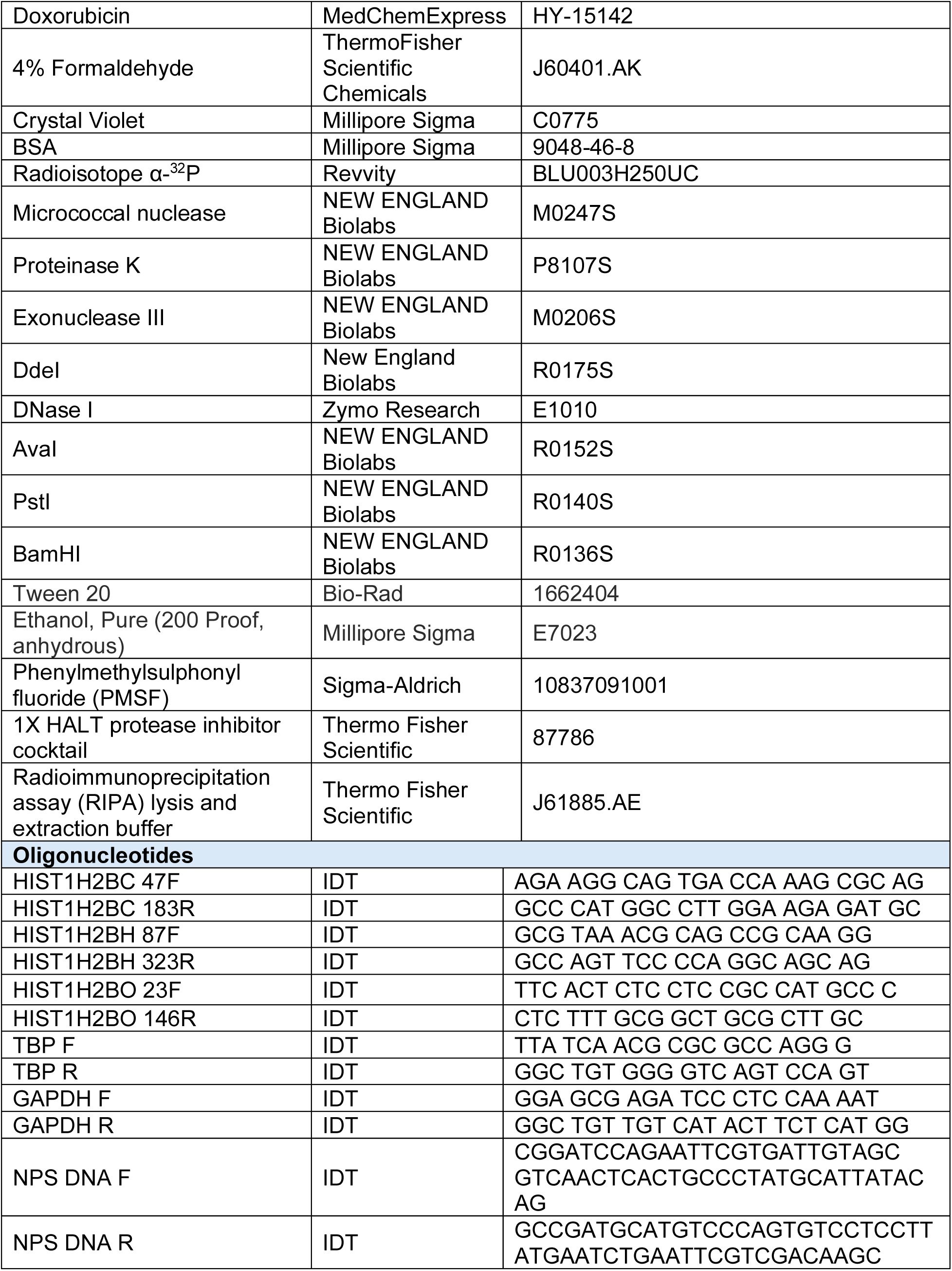

