## Supplemental Figures for "Incorporating histone H2B variants into chromatin modifies chromatin accessibility to induce epithelial-to-mesenchymal transition in breast cancer"

**Fig S1. H2B variant expression in breast cancer subtypes, and other cancers. A**, H2B variant expression in breast tumors (pink) and adjacent normal breast tissue (turquoise), and the corresponding fold-change in gene expression. Data taken from The Cancer Genome Atlas (TCGA). **B**, Principal component analysis (PCA) of TCGA global transcriptomic profiles in breast cancer subtypes. **C**, Certain H2B variants are up-regulated within breast cancer subtypes. Gray dots are those genes with $Log_2$(FoldChange) > 1.2. H2B genes are indicated with red dots. **D**, Kaplan–Meier survival plots where high *HIST1H2BO* gene expression is significantly associated with disease outcomes as indicated by the hazard ratio (HR). Ovarian cancer n=133, p<0.02; colorectal cancer n=62, p<0.05; acute myeloid leukemia (AML) n=163, p<0.004. **E**, Same as panel D, except for *HIST1H2BH* expression. AML n=163, p<0.004; adenocarcinoma n=204, p<0.3; astrocytoma n=77, p<0.002; diffuse large B cell lymphoma (DLBC) n=53, p<0.5.

**Fig S2. Breast cancer patients can be stratified into three clusters based on coordinated H2B variant expression and their co-expressed chromatin modifying genes.** Heatmaps represent unsupervised K-means clustering of patients with the indicated breast cancer subtypes, and are based on the same histone variants identified in **Fig 2A**. Each column is a histone variant gene, and each row is a patient. Data were derived from The Cancer Genome Atlas. Dot plots show significantly enriched C3 oncogenic, Reactome, KEGG, and Biological Process signatures. Fig S2 breaks across multiple pages.

**Fig S3. Population distribution of breast cancer patients from the previously stradified clusters.** Breast cancer patients with high *HIST1H2BO* expression have more Asian and Black/African Americans than that group of patients with low *HIST1H2BO* expression (**A**), and tend to be younger (**B**).

**Fig S4. H2B variants alter nucleosome structure. A**, Reconstituted H2B variant nucleosomes were separated by SDS-PAGE and stained with coomassie blue. All histones are detected, indicating successful reconstitution of intact nucleosomes **B**, Reconstituted H2B variant nucleosomes contained the $^{32}$P-labelled 601 nucleosome positioning sequence (NPS) and free DNA control. **C**, Gel electrophoresis images of H2B variant nucleosomes digested with DNase I for the indicated times. H2B1O nucleosome DNA digests slower, indicating reduced enzyme access to the nucleosomal DNA. **D**, BamHI digestion of free NPS DNA and DNA in the 17-mer nucleosome array. Only the free DNA was digested, indicating successful reconstitution of the nucleosome arrays. **E**, Same as **D**, except digested with AvaI. The band shifts from free DNA to the nucleosome arrays confirm there was no detectable, free NPS DNA within the nucleosome array preparations. **F**, Gel electrophoresis images of H2B variant nucleosome arrays digested with PstI for the indicated times. H2B1O-containing arrays digest slower, indicating reduced enzyme access to the entry/exit middle nucleosomal DNA. Each experiment was repeated four times. **G**, Representative AFM images of nucleosome arrays containing H2B1C, H2B1H, and H2B1O. Each red box is 400 $nm^2$ and is enlarged in the middle column. The right column shows the 3-dimensional topology of chromatin arrays shown in the middle column.

**Fig S5. Transfected H2B variants are stably expressed throughout the cell cycle and incorporated into chromatin. A**, Normalized HA counts from RNA-seq data in MCF10A cell lines expressing the indicated H2B variant minigene. **B**, Flow cytometry scatter plots showing HA-tagged H2B variant expression in G0/1, S, and G2/M cell cycle phases. **C**, Anti-HA immunoblots show that H2B1C, H2B1H, and H2B1O co-localize with H3 in chromatin bound sub-cellular fraction (n=3; biological replicates).

**Fig S6. The top enriched pathways in MCF10A cells expressing *HIST1H2BO*.** Activated inflammatory pathways are indicated by red text.

**Fig S7. Top enriched genes and pathways in MCF10A cells expressing (A) *HIST1H2BC* and (B) *HISTH2BH*.** Fig S7 breaks across multiple pages.

**Fig S8. RNA-seq analysis of cytokine-transformed MCF10A cells.** The top 30 differentially expressed genes (DEGs) relative to an empty vector (EV) control. Dot plots illustrate the top enriched signatures.

**Fig S9. Integrated ATAC-seq and RNA-seq. A**, Overlap between opened chromatin regions (ATAC UP) and upregulated gene expression (RNA UP), and closed chromatin regions (ATAC DN) and down-regulated gene expression (RNA DN). Counts represent the number of genes with >1 $Log_2$(FoldChange) in gene expression, and >1 shrunken Log(FoldChange) for open chromatin. **B-D**, Over-representation analysis of the most enriched (**B**) C6 oncogenic, (**C**) Hallmark, and (**D**) biological process signatures where ATAC-seq and RNA-seq data overlapped. **E-F**, Enrichment maps of the RAS-RAF-MER pathway and overlapping genes in cytokine-transformed cells and cells expressing (**E**) *HIST1H2BC* and (**F**) *HIST1H2BH*. Cells transfected with an empty vector (EV) served as the control. **G**, Top enriched C6 oncogenic and Hallmark signatures in cytokine-transformed cells that were previously transfected with H2B minigenes. H21BO Hallmark signatures are in Fig 8B. Fig S9 breaks across multiple pages.

**Fig S1**

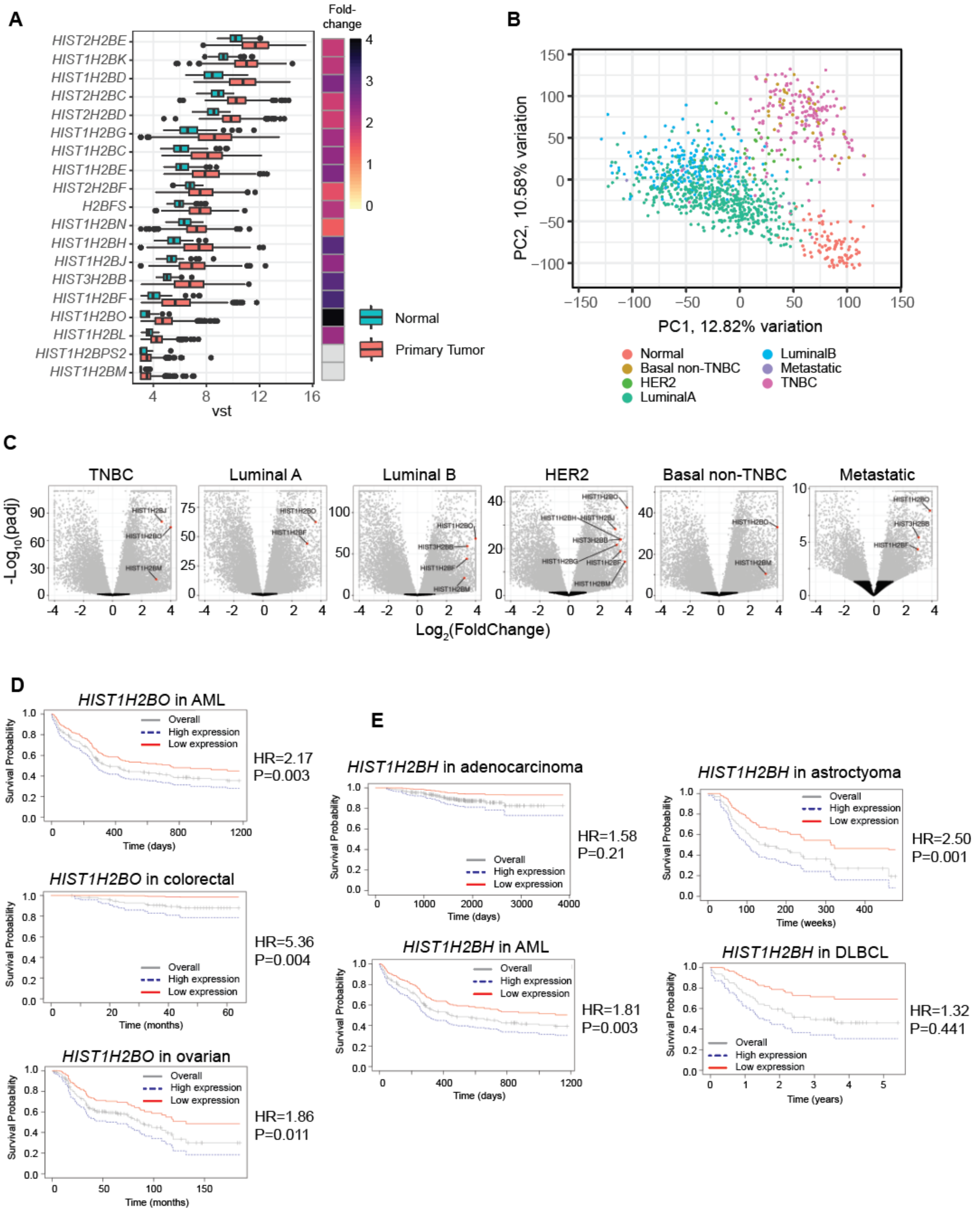

A
HIST2H2BE
HIST1H2BK
HIST1H2BD
HIST2H2BC
HIST2H2BD
HIST1H2BG
HIST1H2BC
HIST1H2BE
HIST2H2BF
H2BFS
HIST1H2BN
HIST1H2BH
HIST1H2BJ
HIST3H2BB
HIST1H2BF
HIST1H2BO
HIST1H2BL
HIST1H2BPS2
HIST1H2BM
vst
Fold-change
Normal
Primary Tumor
B
PC1, 12.82% variation
PC2, 10.58% variation
Normal
Basal non-TNBC
HER2
LuminalA
LuminalB
Metastatic
TNBC
C
TNBC
Luminal A
Luminal B
HER2
Basal non-TNBC
Metastatic
-Log10(padj)
Log2(FoldChange)
D
HIST1H2BO in AML
HR=2.17
P=0.003
HIST1H2BO in colorectal
HR=5.36
P=0.004
HIST1H2BO in ovarian
HR=1.86
P=0.011
E
HIST1H2BH in adenocarcinoma
HR=1.58
P=0.21
HIST1H2BH in astroctyoma
HR=2.50
P=0.001
HIST1H2BH in AML
HR=1.81
P=0.003
HIST1H2BH in DLBCL
HR=1.32
P=0.441
Survival Probability
Overall
High expression
Low expression
Time (days)
Time (months)
Time (weeks)
Time (years)

**Fig S2**

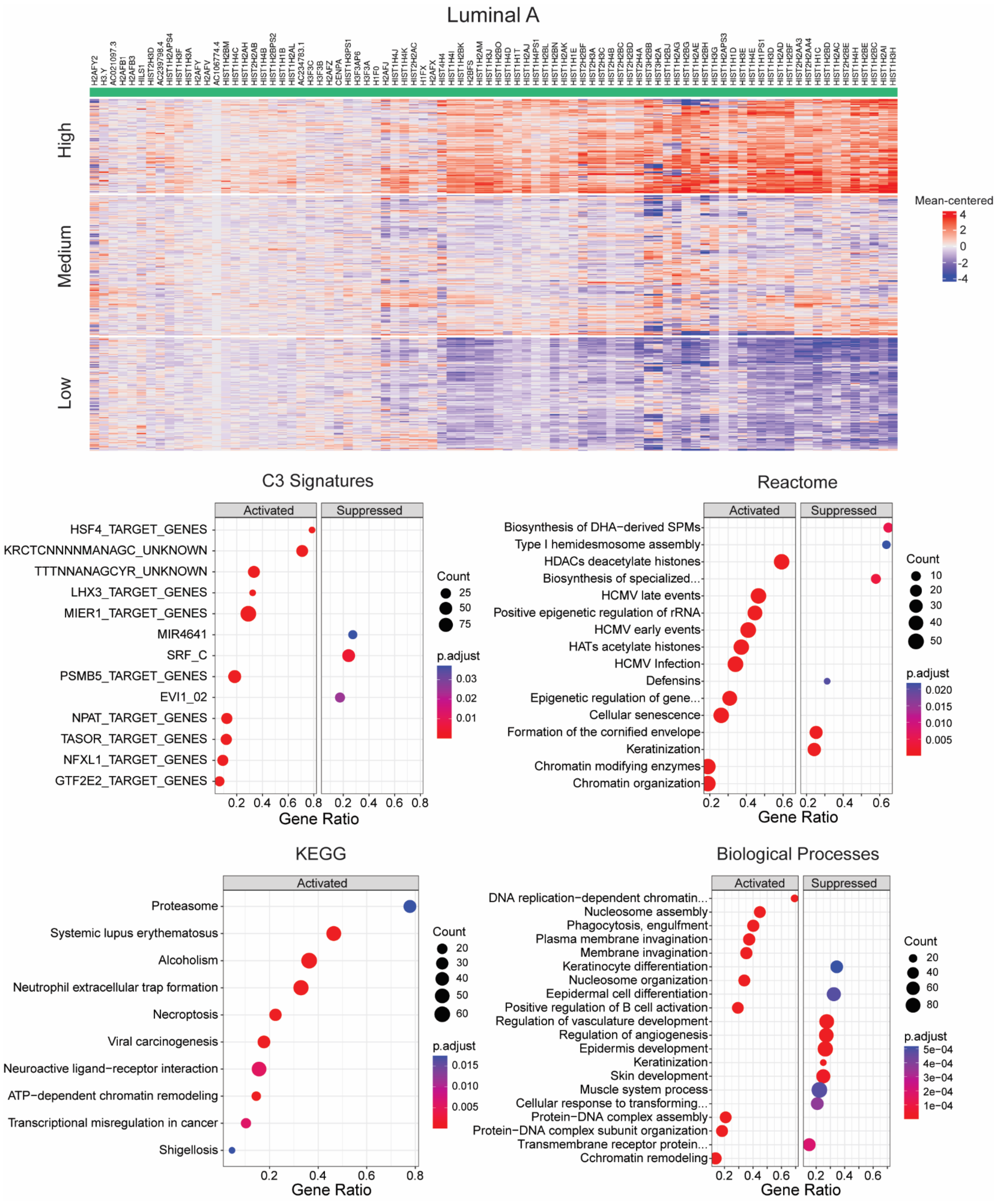

Luminal A
High
Medium
Low
Mean-centered
4
2
0
-2
-4
C3 Signatures
Activated
Suppressed
HSF4_TARGET_GENES
KRCTCNNNNMANAGC_UNKNOWN
TTTNNANAGCYR_UNKNOWN
LHX3_TARGET_GENES
MIER1_TARGET_GENES
MIR4641
SRF_C
PSMB5_TARGET_GENES
EVI1_02
NPAT_TARGET_GENES
TASOR_TARGET_GENES
NFXL1_TARGET_GENES
GTF2E2_TARGET_GENES
Count
25
50
75
p.adjust
0.03
0.02
0.01
Gene Ratio
Reactome
Activated
Suppressed
Biosynthesis of DHA−derived SPMs
Type I hemidesmosome assembly
HDACs deacetylate histones
Biosynthesis of specialized...
HCMV late events
Positive epigenetic regulation of rRNA
HCMV early events
HATs acetylate histones
HCMV Infection
Defensins
Epigenetic regulation of gene...
Cellular senescence
Formation of the cornified envelope
Keratinization
Chromatin modifying enzymes
Chromatin organization
Count
10
20
30
40
50
p.adjust
0.020
0.015
0.010
0.005
Gene Ratio
KEGG
Activated
Proteasome
Systemic lupus erythematosus
Alcoholism
Neutrophil extracellular trap formation
Necroptosis
Viral carcinogenesis
Neuroactive ligand−receptor interaction
ATP−dependent chromatin remodeling
Transcriptional misregulation in cancer
Shigellosis
Count
20
30
40
50
60
p.adjust
0.015
0.010
0.005
Gene Ratio
Biological Processes
Activated
Suppressed
DNA replication−dependent chromatin...
Nucleosome assembly
Phagocytosis, engulfment
Plasma membrane invagination
Membrane invagination
Keratinocyte differentiation
Nucleosome organization
Epidermal cell differentiation
Positive regulation of B cell activation
Regulation of vasculature development
Regulation of angiogenesis
Epidermis development
Keratinization
Skin development
Muscle system process
Cellular response to transforming...
Protein−DNA complex assembly
Protein−DNA complex subunit organization
Transmembrane receptor protein...
Cchromatin remodeling
Count
20
40
60
80
p.adjust
5e−04
4e−04
3e−04
2e−04
1e−04
Gene Ratio

Fig S2, continued

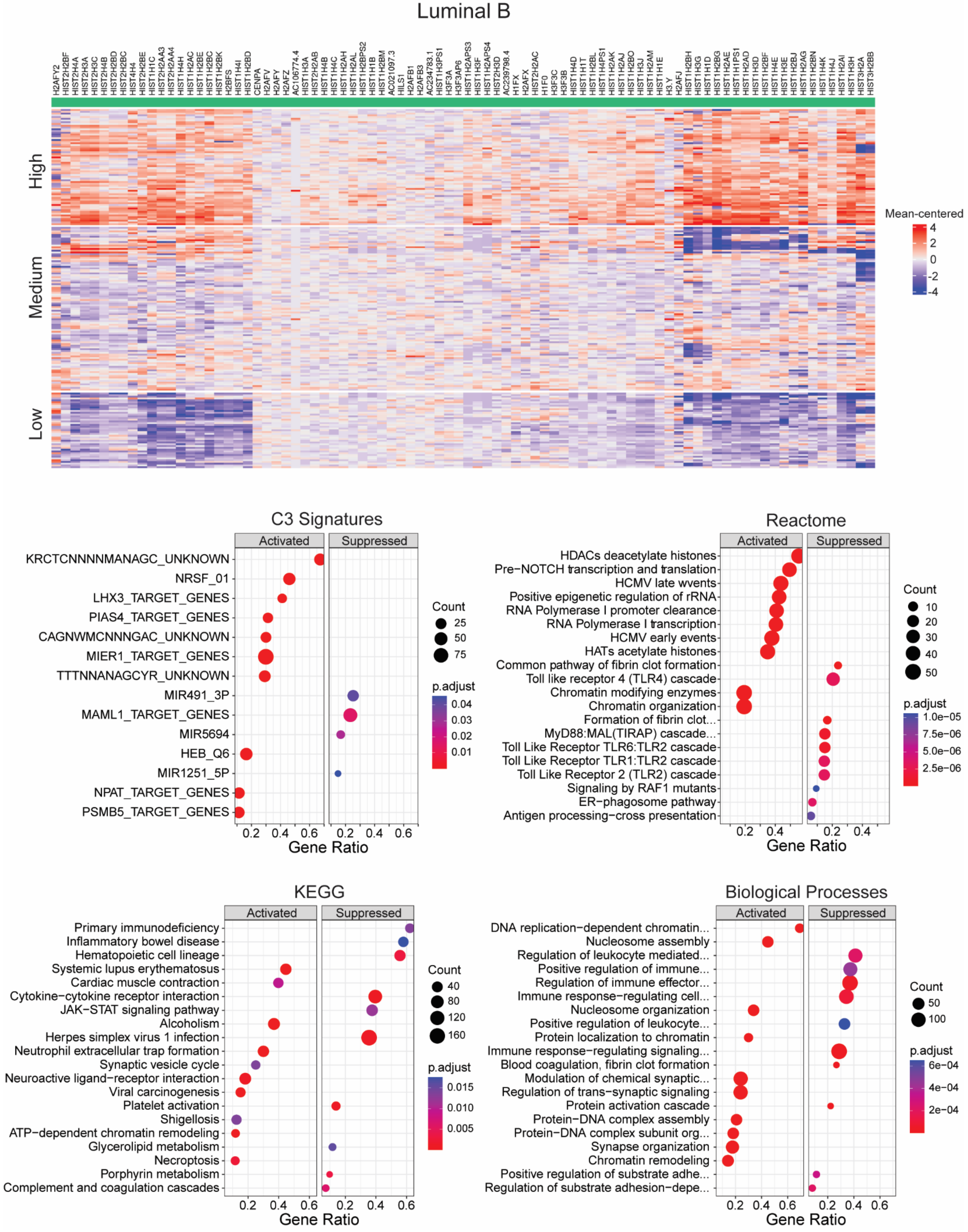

Fig S2, continued

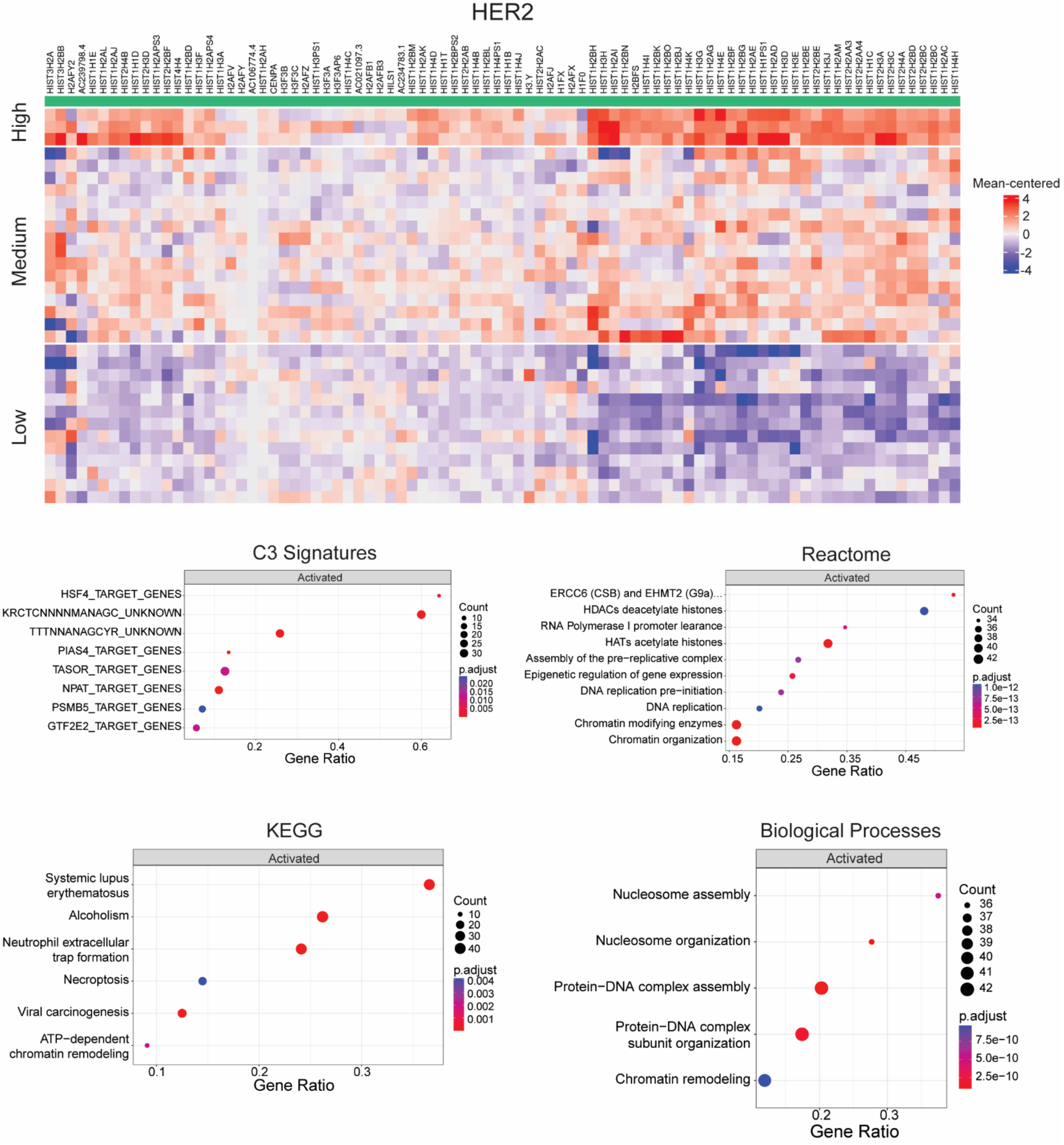

Fig S2, continued

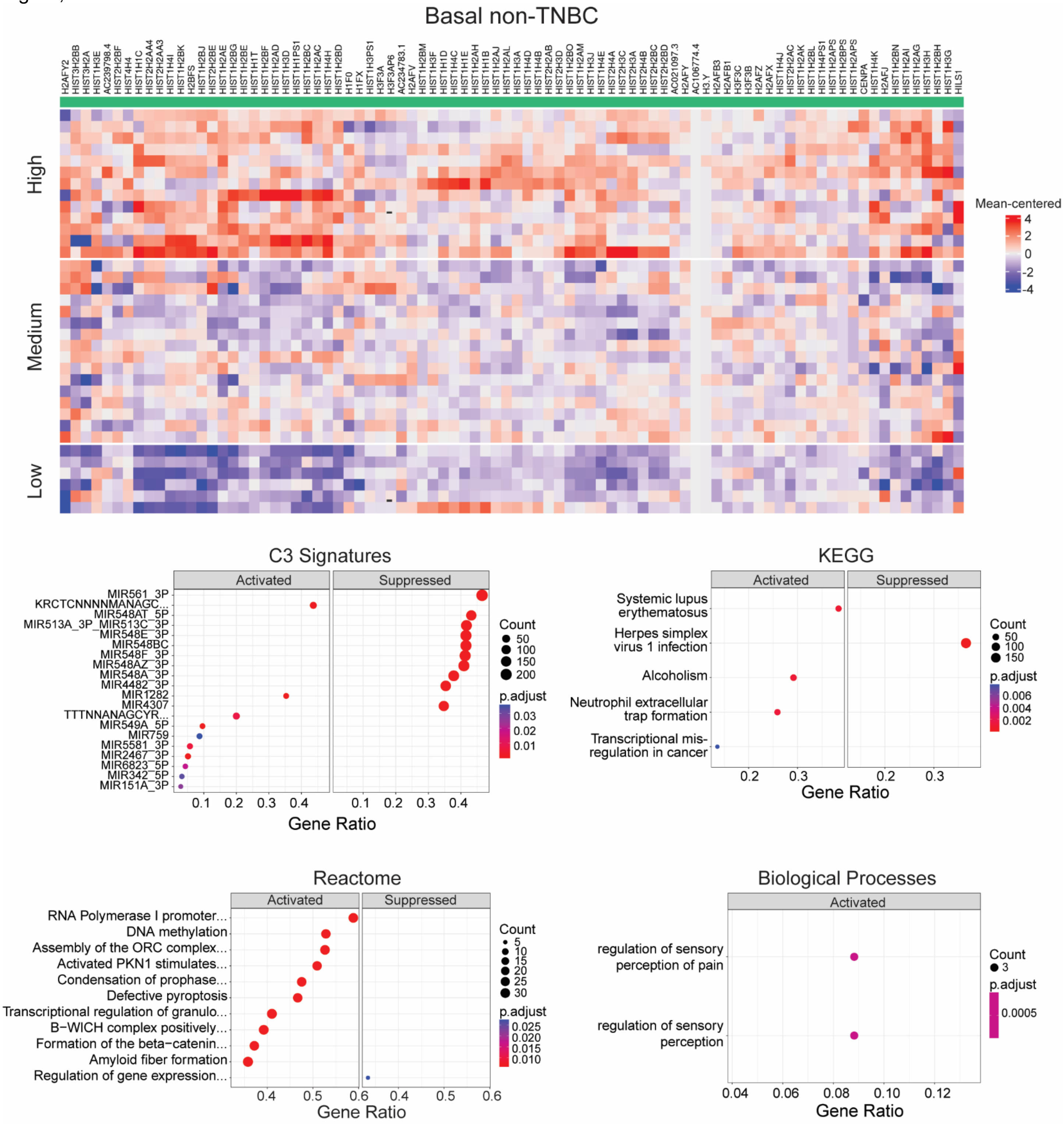

Fig S2, continued

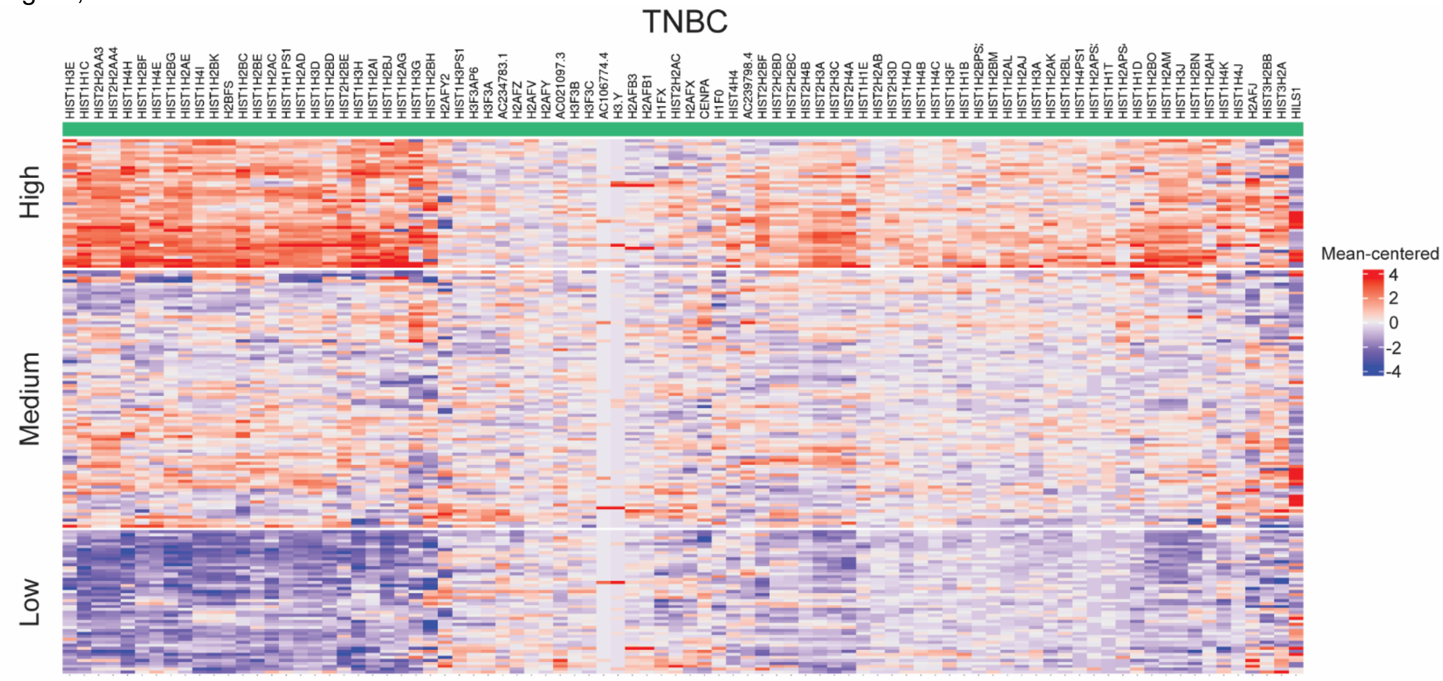


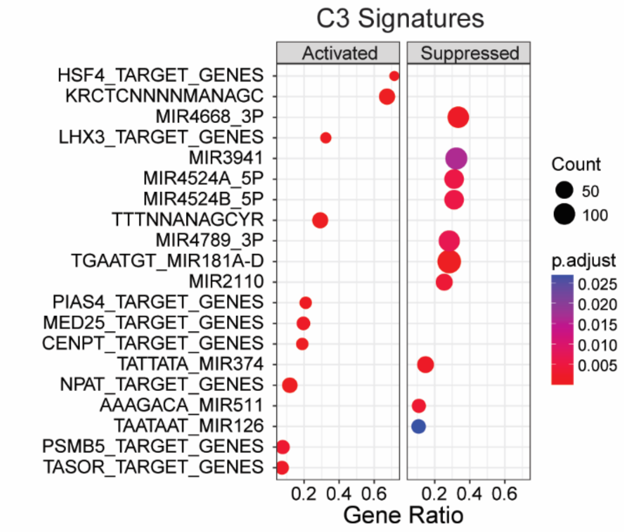


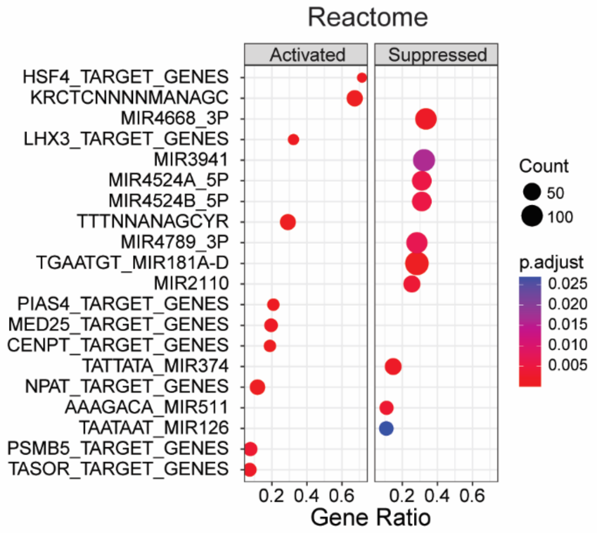


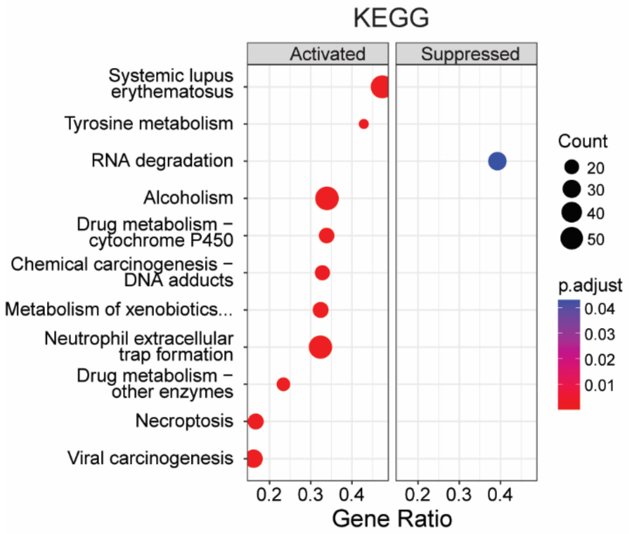


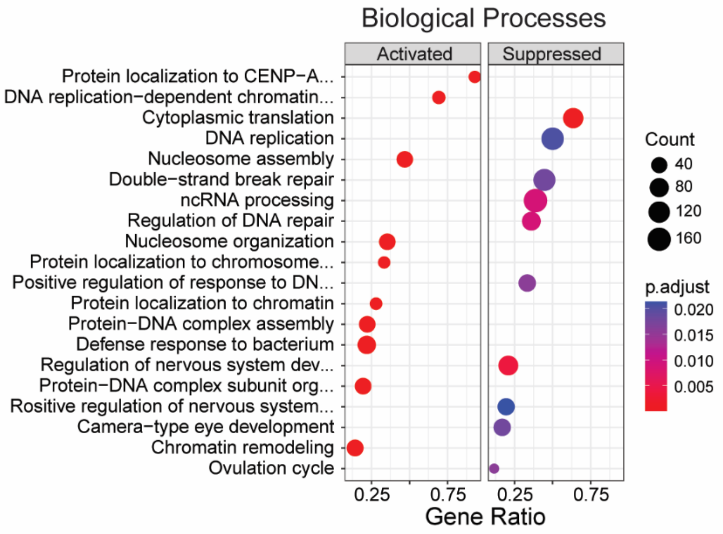

Fig S3

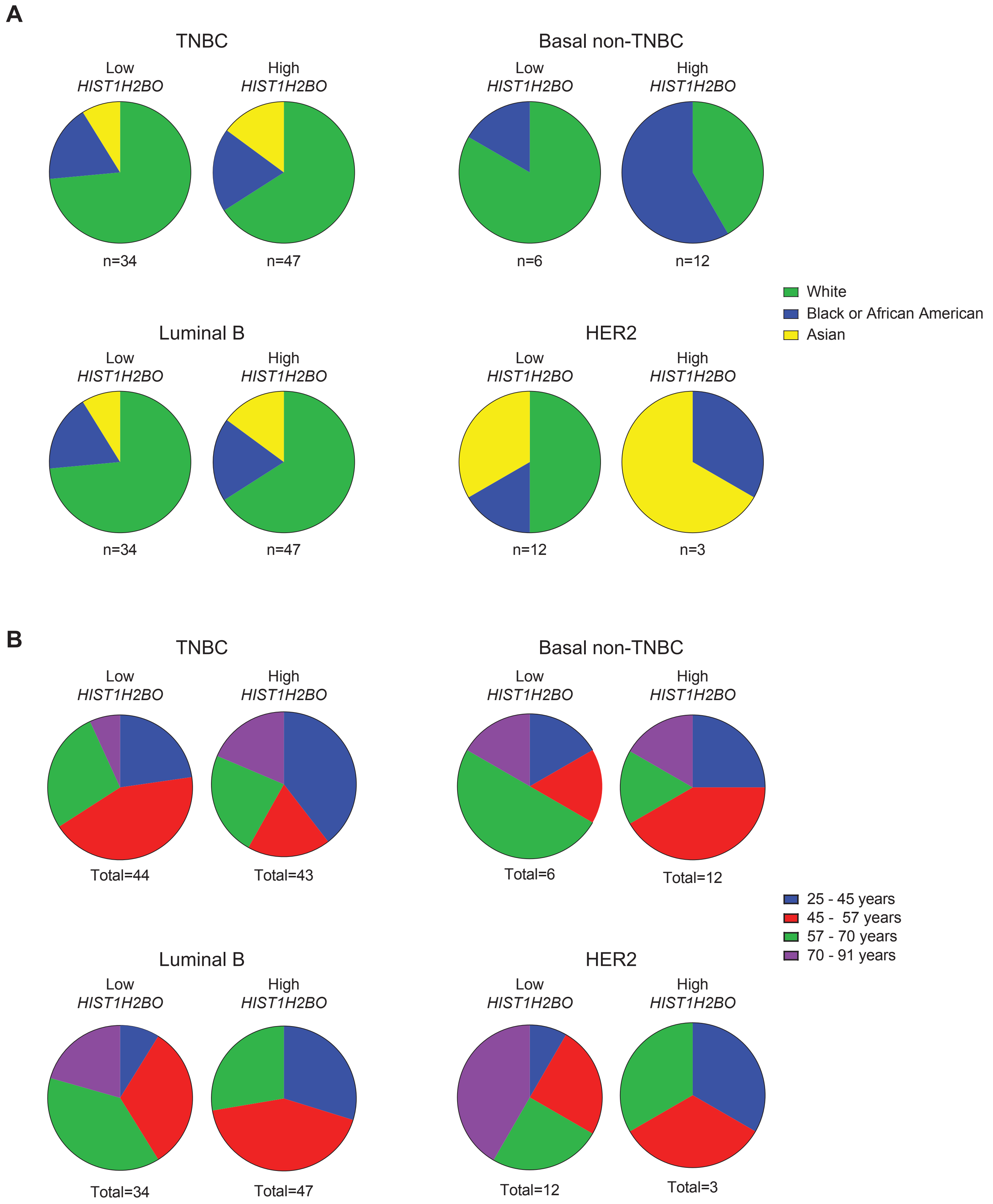
A
TNBC
Low
HIST1H2BO
High
HIST1H2BO
n=34
n=47
Basal non-TNBC
Low
HIST1H2BO
High
HIST1H2BO
n=6
n=12
White
Black or African American
Asian
Luminal B
Low
HIST1H2BO
High
HIST1H2BO
n=34
n=47
HER2
Low
HIST1H2BO
High
HIST1H2BO
n=12
n=3
B
TNBC
Low
HIST1H2BO
High
HIST1H2BO
Total=44
Total=43
Basal non-TNBC
Low
HIST1H2BO
High
HIST1H2BO
Total=6
Total=12
25 - 45 years
45 - 57 years
57 - 70 years
70 - 91 years
Luminal B
Low
HIST1H2BO
High
HIST1H2BO
Total=34
Total=47
HER2
Low
HIST1H2BO
High
HIST1H2BO
Total=12
Total=3

Fig S4

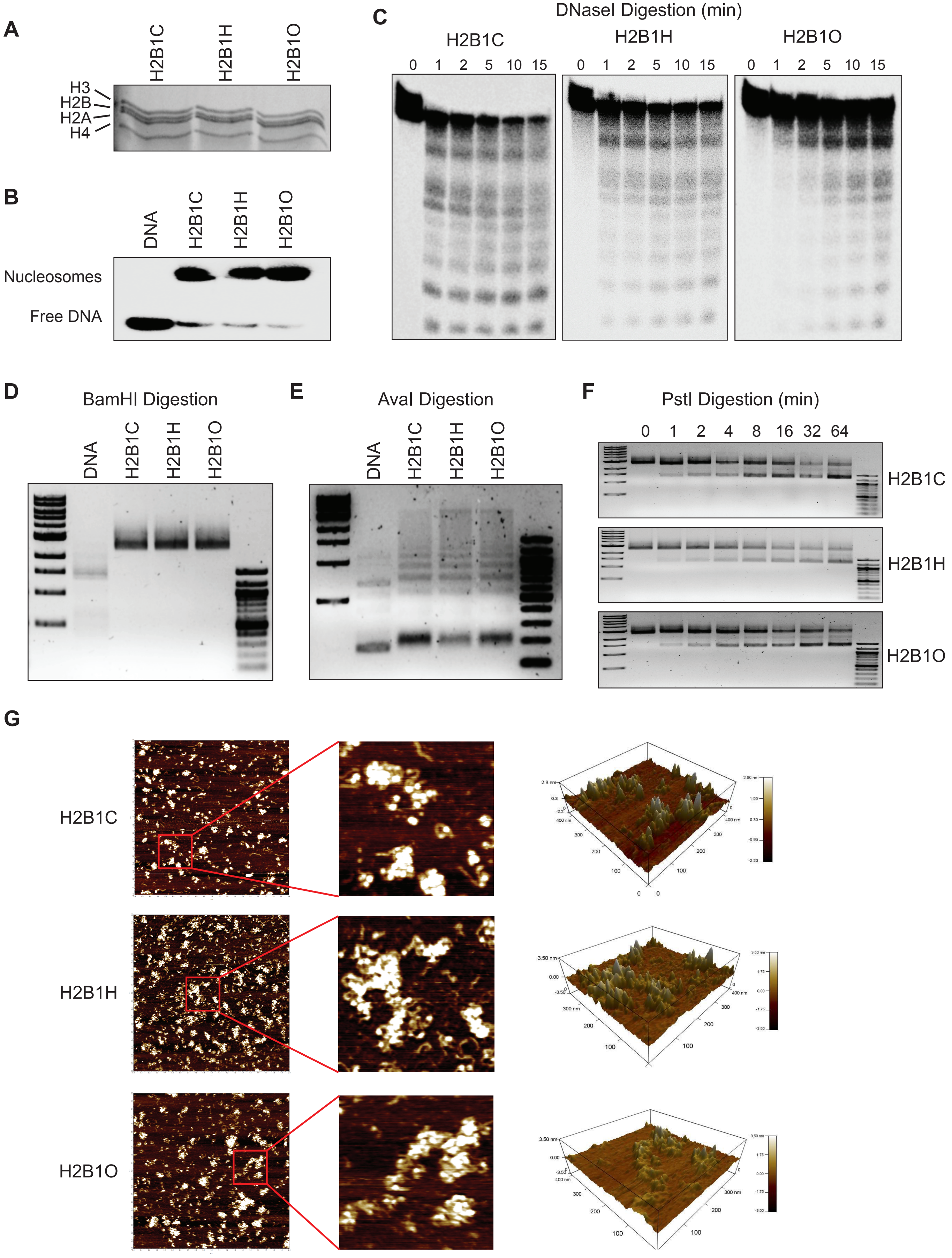
A
H2B1C
H2B1H
H2B1O
H3
H2B
H2A
H4
B
DNA
H2B1C
H2B1H
H2B1O
Nucleosomes
Free DNA
C
DNaseI Digestion (min)
H2B1C
H2B1H
H2B1O
0 1 2 5 10 15
0 1 2 5 10 15
0 1 2 5 10 15
D
BamHI Digestion
DNA
H2B1C
H2B1H
H2B1O
E
AvaI Digestion
DNA
H2B1C
H2B1H
H2B1O
F
PstI Digestion (min)
0 1 2 4 8 16 32 64
H2B1C
H2B1H
H2B1O
G
H2B1C
H2B1H
H2B1O

Fig S5

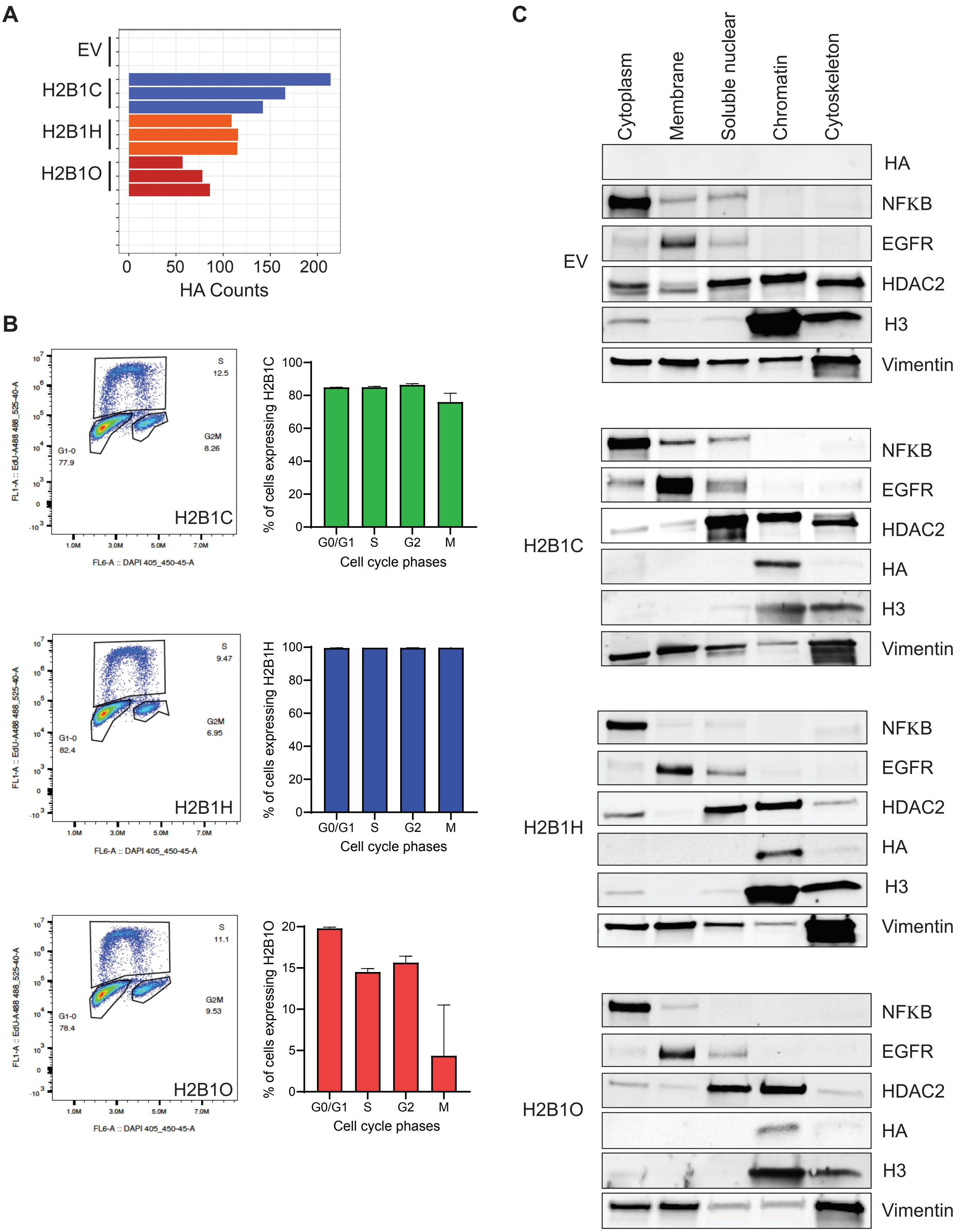
A
EV
H2B1C
H2B1H
H2B1O
0
50
100
150
200
HA Counts
B
S
12.5
G2M
8.26
G1-0
77.9
H2B1C
FL6-A :: DAPI 405_450-45-A
FL1-A :: EdU-A488 488_525-40-A
% of cells expressing H2B1C
G0/G1
S
G2
M
Cell cycle phases
S
9.47
G2M
6.95
G1-0
82.4
H2B1H
% of cells expressing H2B1H
S
11.1
G2M
9.53
G1-0
78.4
H2B1O
% of cells expressing H2B1O
C
Cytoplasm
Membrane
Soluble nuclear
Chromatin
Cytoskeleton
EV
HA
NFKB
EGFR
HDAC2
H3
Vimentin
H2B1C
H2B1H
H2B1O

**Fig S6**

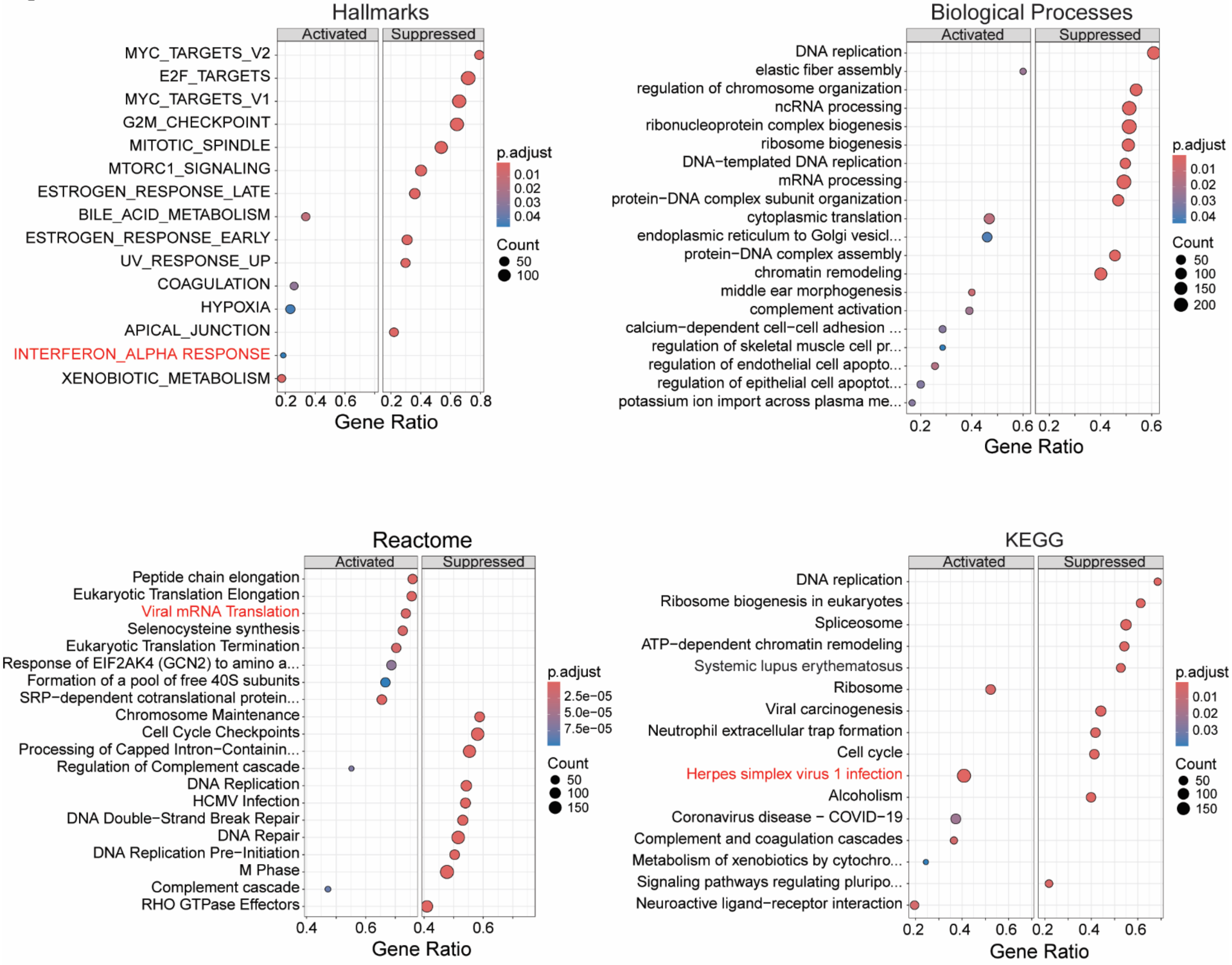
Hallmarks
Activated
Suppressed
MYC_TARGETS_V2
E2F_TARGETS
MYC_TARGETS_V1
G2M_CHECKPOINT
MITOTIC_SPINDLE
MTORC1_SIGNALING
ESTROGEN_RESPONSE_LATE
BILE_ACID_METABOLISM
ESTROGEN_RESPONSE_EARLY
UV_RESPONSE_UP
COAGULATION
HYPOXIA
APICAL_JUNCTION
INTERFERON_ALPHA RESPONSE
XENOBIOTIC_METABOLISM
Gene Ratio
p.adjust
Count
Biological Processes
DNA replication
elastic fiber assembly
regulation of chromosome organization
ncRNA processing
ribonucleoprotein complex biogenesis
ribosome biogenesis
DNA−templated DNA replication
mRNA processing
protein−DNA complex subunit organization
cytoplasmic translation
endoplasmic reticulum to Golgi vesicl...
protein−DNA complex assembly
chromatin remodeling
middle ear morphogenesis
complement activation
calcium−dependent cell−cell adhesion ...
regulation of skeletal muscle cell pr...
regulation of endothelial cell apopto...
regulation of epithelial cell apoptot...
potassium ion import across plasma me...
Reactome
Peptide chain elongation
Eukaryotic Translation Elongation
Viral mRNA Translation
Selenocysteine synthesis
Eukaryotic Translation Termination
Response of EIF2AK4 (GCN2) to amino a...
Formation of a pool of free 40S subunits
SRP−dependent cotranslational protein...
Chromosome Maintenance
Cell Cycle Checkpoints
Processing of Capped Intron−Containin...
Regulation of Complement cascade
DNA Replication
HCMV Infection
DNA Double−Strand Break Repair
DNA Repair
DNA Replication Pre−Initiation
M Phase
Complement cascade
RHO GTPase Effectors
KEGG
DNA replication
Ribosome biogenesis in eukaryotes
Spliceosome
ATP−dependent chromatin remodeling
Systemic lupus erythematosus
Ribosome
Viral carcinogenesis
Neutrophil extracellular trap formation
Cell cycle
Herpes simplex virus 1 infection
Alcoholism
Coronavirus disease − COVID−19
Complement and coagulation cascades
Metabolism of xenobiotics by cytochro...
Signaling pathways regulating pluripo...
Neuroactive ligand−receptor interaction

**Fig S7**

**A**

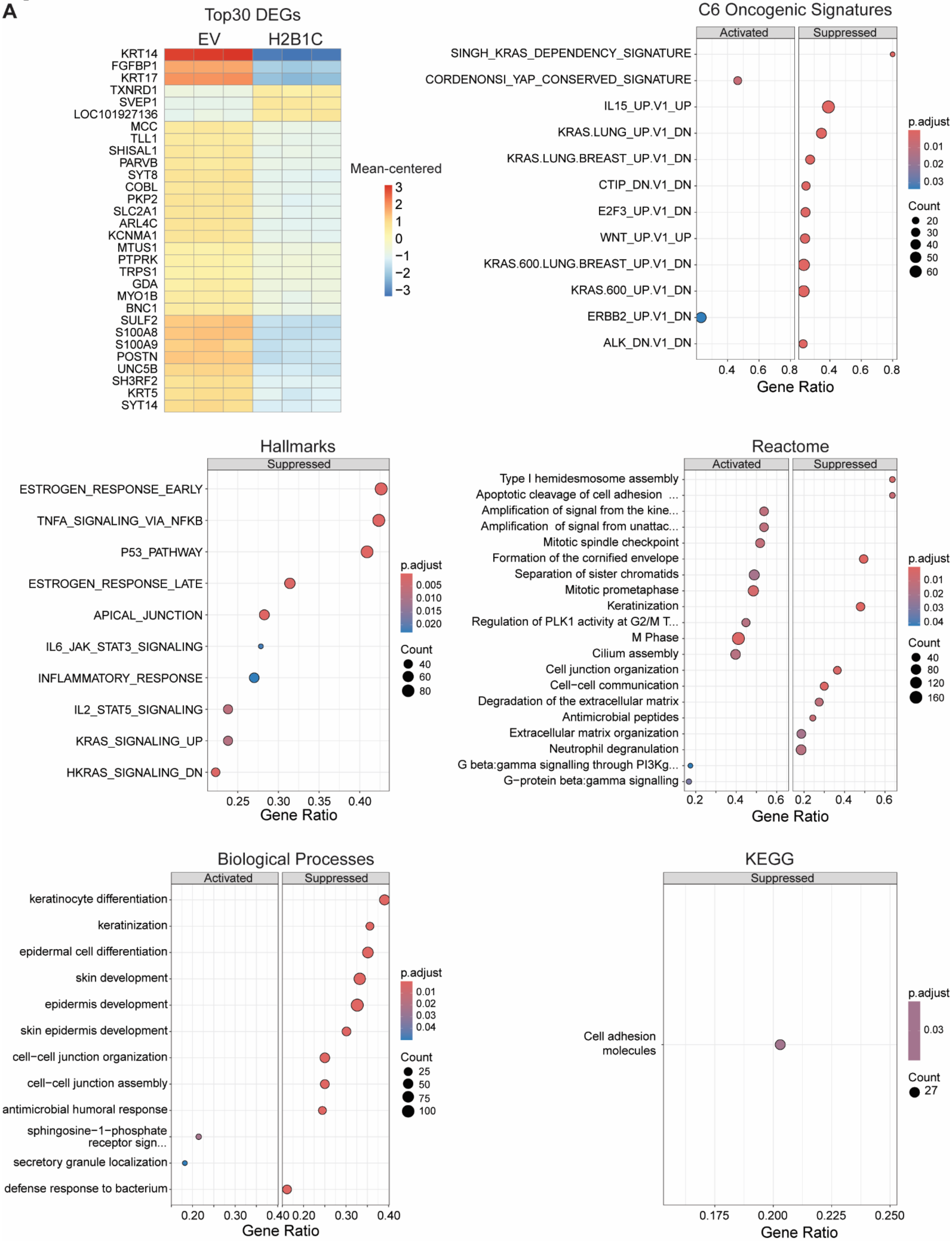
Top30 DEGs
EV
H2B1C
KRT14
FGFBP1
KRT17
TXNRD1
SVEP1
LOC101927136
MCC
TLL1
SHISAL1
PARVB
SYT8
COBL
PKP2
SLC2A1
ARL4C
KCNMA1
MTUS1
PTPRK
TRPS1
GDA
MYO1B
BNC1
SULF2
S100A8
S100A9
POSTN
UNC5B
SH3RF2
KRT5
SYT14
Mean-centered
3
2
1
0
−1
−2
−3
C6 Oncogenic Signatures
Activated
Suppressed
SINGH_KRAS_DEPENDENCY_SIGNATURE
CORDENONSI_YAP_CONSERVED_SIGNATURE
IL15_UP.V1_UP
KRAS.LUNG_UP.V1_DN
KRAS.LUNG.BREAST_UP.V1_DN
CTIP_DN.V1_DN
E2F3_UP.V1_DN
WNT_UP.V1_UP
KRAS.600.LUNG.BREAST_UP.V1_DN
KRAS.600_UP.V1_DN
ERBB2_UP.V1_DN
ALK_DN.V1_DN
0.4 0.6 0.8 0.4 0.6 0.8
Gene Ratio
p.adjust
0.01
0.02
0.03
Count
20
30
40
50
60
Hallmarks
Suppressed
ESTROGEN_RESPONSE_EARLY
TNFA_SIGNALING_VIA_NFKB
P53_PATHWAY
ESTROGEN_RESPONSE_LATE
APICAL_JUNCTION
IL6_JAK_STAT3_SIGNALING
INFLAMMATORY_RESPONSE
IL2_STAT5_SIGNALING
KRAS_SIGNALING_UP
HKRAS_SIGNALING_DN
0.25 0.30 0.35 0.40
Gene Ratio
p.adjust
0.005
0.010
0.015
0.020
Count
40
60
80
Reactome
Activated
Suppressed
Type I hemidesmosome assembly
Apoptotic cleavage of cell adhesion ...
Amplification of signal from the kine...
Amplification of signal from unattac...
Mitotic spindle checkpoint
Formation of the cornified envelope
Separation of sister chromatids
Mitotic prometaphase
Keratinization
Regulation of PLK1 activity at G2/M T...
M Phase
Cilium assembly
Cell junction organization
Cell−cell communication
Degradation of the extracellular matrix
Antimicrobial peptides
Extracellular matrix organization
Neutrophil degranulation
G beta:gamma signalling through PI3Kg...
G−protein beta:gamma signalling
0.2 0.4 0.6 0.2 0.4 0.6
Gene Ratio
p.adjust
0.01
0.02
0.03
0.04
Count
40
80
120
160
Biological Processes
Activated
Suppressed
keratinocyte differentiation
keratinization
epidermal cell differentiation
skin development
epidermis development
skin epidermis development
cell−cell junction organization
cell−cell junction assembly
antimicrobial humoral response
sphingosine−1−phosphate receptor sign...
secretory granule localization
defense response to bacterium
0.20 0.30 0.40 0.20 0.30 0.40
Gene Ratio
p.adjust
0.01
0.02
0.03
0.04
Count
25
50
75
100
KEGG
Suppressed
Cell adhesion molecules
0.175 0.200 0.225 0.250
Gene Ratio
p.adjust
0.03
Count
27

Fig 7, continued

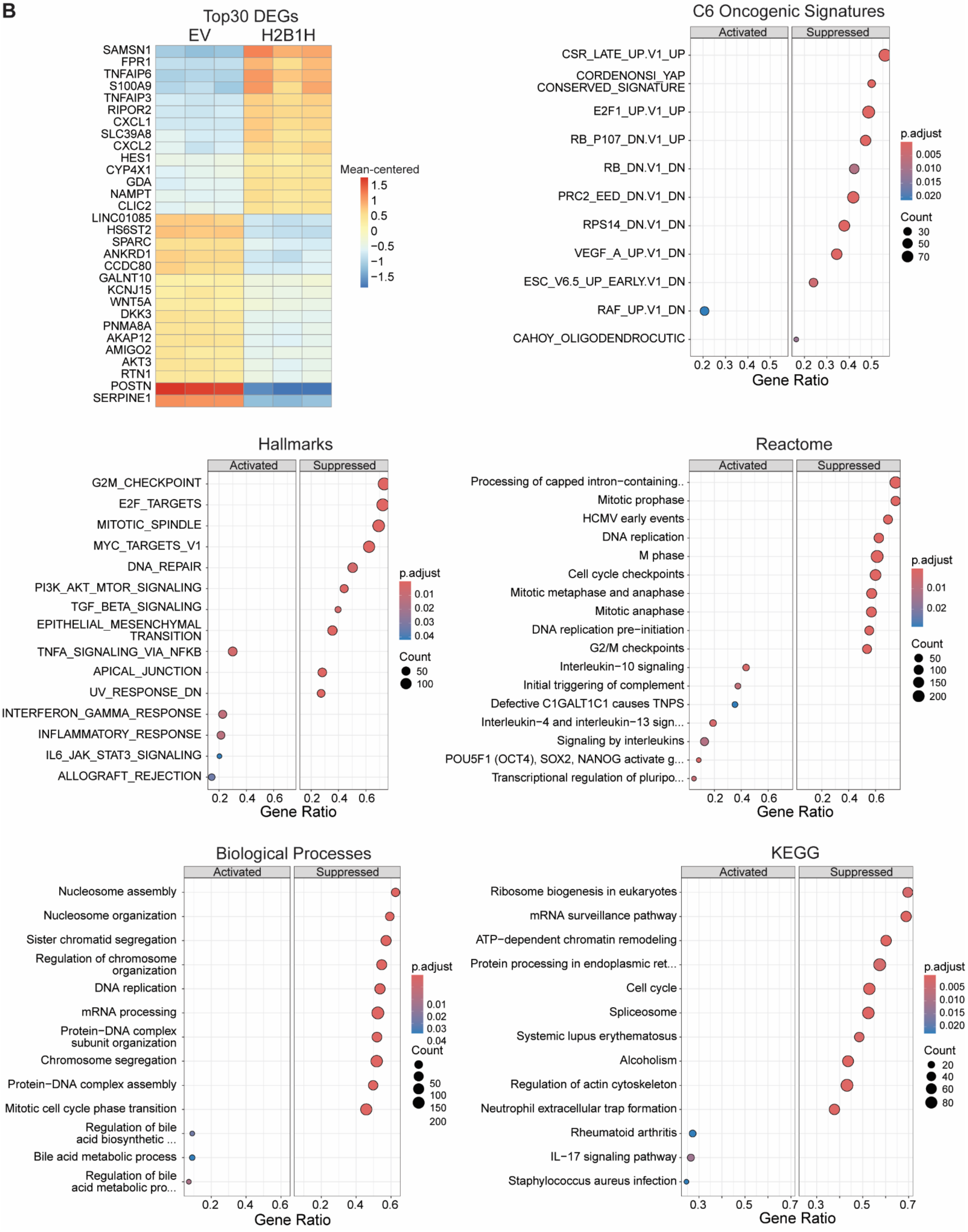

**Fig S8**

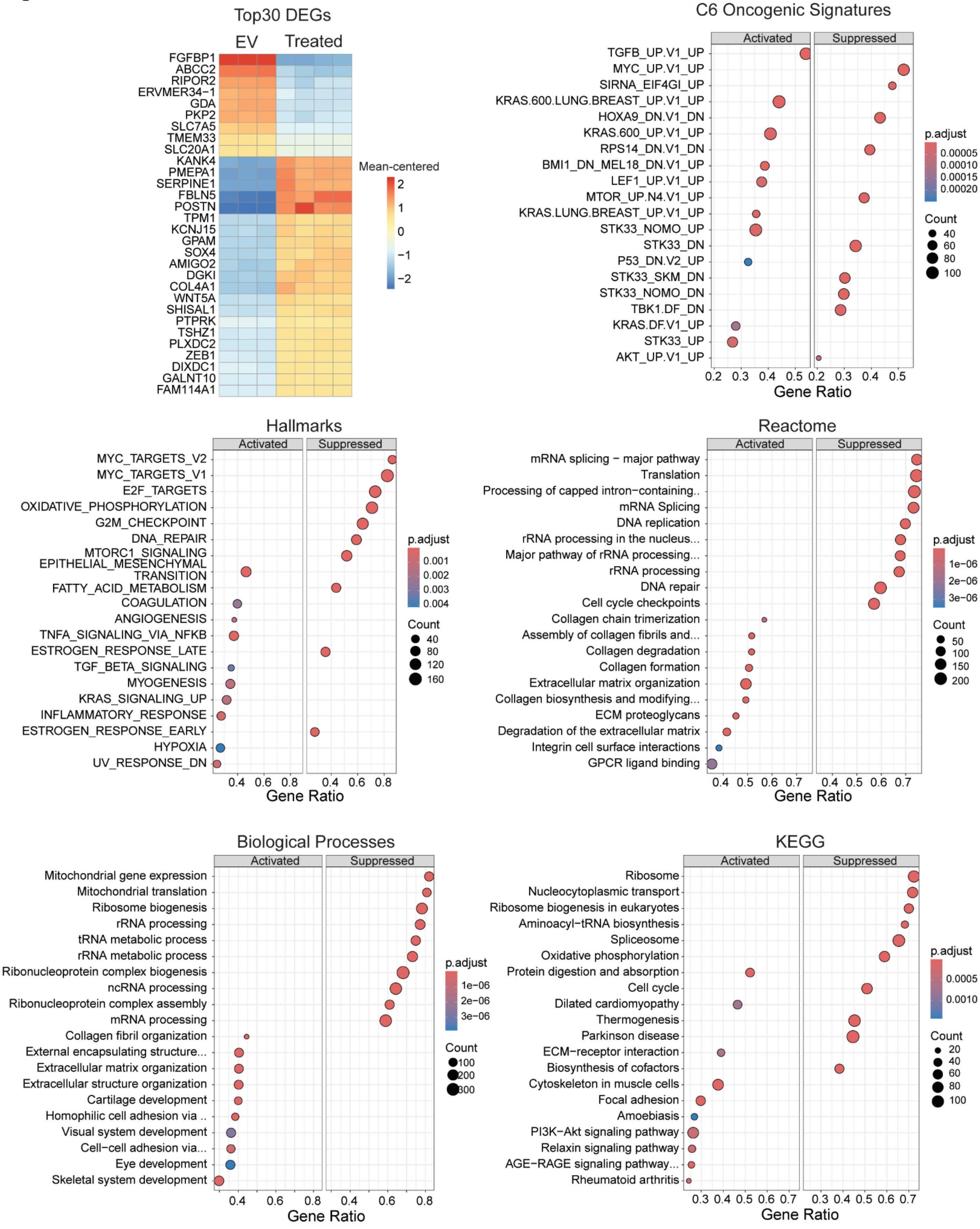

Top30 DEGs
EV
Treated
FGFBP1
ABCC2
RIPOR2
ERVMER34−1
GDA
PKP2
SLC7A5
TMEM33
SLC20A1
KANK4
PMEPA1
SERPINE1
FBLN5
POSTN
TPM1
KCNJ15
GPAM
SOX4
AMIGO2
DGKI
COL4A1
WNT5A
SHISAL1
PTPRK
TSHZ1
PLXDC2
ZEB1
DIXDC1
GALNT10
FAM114A1
Mean-centered
2
1
0
−1
−2
C6 Oncogenic Signatures
Activated
Suppressed
TGFB_UP.V1_UP
MYC_UP.V1_UP
SIRNA_EIF4GI_UP
KRAS.600.LUNG.BREAST_UP.V1_UP
HOXA9_DN.V1_DN
KRAS.600_UP.V1_UP
RPS14_DN.V1_DN
BMI1_DN_MEL18_DN.V1_UP
LEF1_UP.V1_UP
MTOR_UP.N4.V1_UP
KRAS.LUNG.BREAST_UP.V1_UP
STK33_NOMO_UP
STK33_DN
P53_DN.V2_UP
STK33_SKM_DN
STK33_NOMO_DN
TBK1.DF_DN
KRAS.DF.V1_UP
STK33_UP
AKT_UP.V1_UP
0.2 0.3 0.4 0.5 0.2 0.3 0.4 0.5
Gene Ratio
p.adjust
0.00005
0.00010
0.00015
0.00020
Count
40
60
80
100
Hallmarks
Activated
Suppressed
MYC_TARGETS_V2
MYC_TARGETS_V1
E2F_TARGETS
OXIDATIVE_PHOSPHORYLATION
G2M_CHECKPOINT
DNA_REPAIR
MTORC1_SIGNALING
EPITHELIAL_MESENCHYMAL TRANSITION
FATTY_ACID_METABOLISM
COAGULATION
ANGIOGENESIS
TNFA_SIGNALING_VIA_NFKB
ESTROGEN_RESPONSE_LATE
TGF_BETA_SIGNALING
MYOGENESIS
KRAS_SIGNALING_UP
INFLAMMATORY_RESPONSE
ESTROGEN_RESPONSE_EARLY
HYPOXIA
UV_RESPONSE_DN
0.4 0.6 0.8 0.4 0.6 0.8
Gene Ratio
p.adjust
0.001
0.002
0.003
0.004
Count
40
80
120
160
Reactome
Activated
Suppressed
mRNA splicing − major pathway
Translation
Processing of capped intron−containing..
mRNA Splicing
DNA replication
rRNA processing in the nucleus...
Major pathway of rRNA processing...
rRNA processing
DNA repair
Cell cycle checkpoints
Collagen chain trimerization
Assembly of collagen fibrils and...
Collagen degradation
Collagen formation
Extracellular matrix organization
Collagen biosynthesis and modifying...
ECM proteoglycans
Degradation of the extracellular matrix
Integrin cell surface interactions
GPCR ligand binding
0.4 0.5 0.6 0.7 0.4 0.5 0.6 0.7
Gene Ratio
p.adjust
1e−06
2e−06
3e−06
Count
50
100
150
200
Biological Processes
Activated
Suppressed
Mitochondrial gene expression
Mitochondrial translation
Ribosome biogenesis
rRNA processing
tRNA metabolic process
rRNA metabolic process
Ribonucleoprotein complex biogenesis
ncRNA processing
Ribonucleoprotein complex assembly
mRNA processing
Collagen fibril organization
External encapsulating structure...
Extracellular matrix organization
Extracellular structure organization
Cartilage development
Homophilic cell adhesion via ..
Visual system development
Cell−cell adhesion via...
Eye development
Skeletal system development
0.4 0.6 0.8 0.4 0.6 0.8
Gene Ratio
p.adjust
1e−06
2e−06
3e−06
Count
100
200
300
KEGG
Activated
Suppressed
Ribosome
Nucleocytoplasmic transport
Ribosome biogenesis in eukaryotes
Aminoacyl−tRNA biosynthesis
Spliceosome
Oxidative phosphorylation
Protein digestion and absorption
Cell cycle
Dilated cardiomyopathy
Thermogenesis
Parkinson disease
ECM−receptor interaction
Biosynthesis of cofactors
Cytoskeleton in muscle cells
Focal adhesion
Amoebiasis
PI3K−Akt signaling pathway
Relaxin signaling pathway
AGE−RAGE signaling pathway...
Rheumatoid arthritis
0.3 0.4 0.5 0.6 0.7 0.3 0.4 0.5 0.6 0.7
Gene Ratio
p.adjust
0.0005
0.0010
Count
20
40
60
80
100

Fig S9

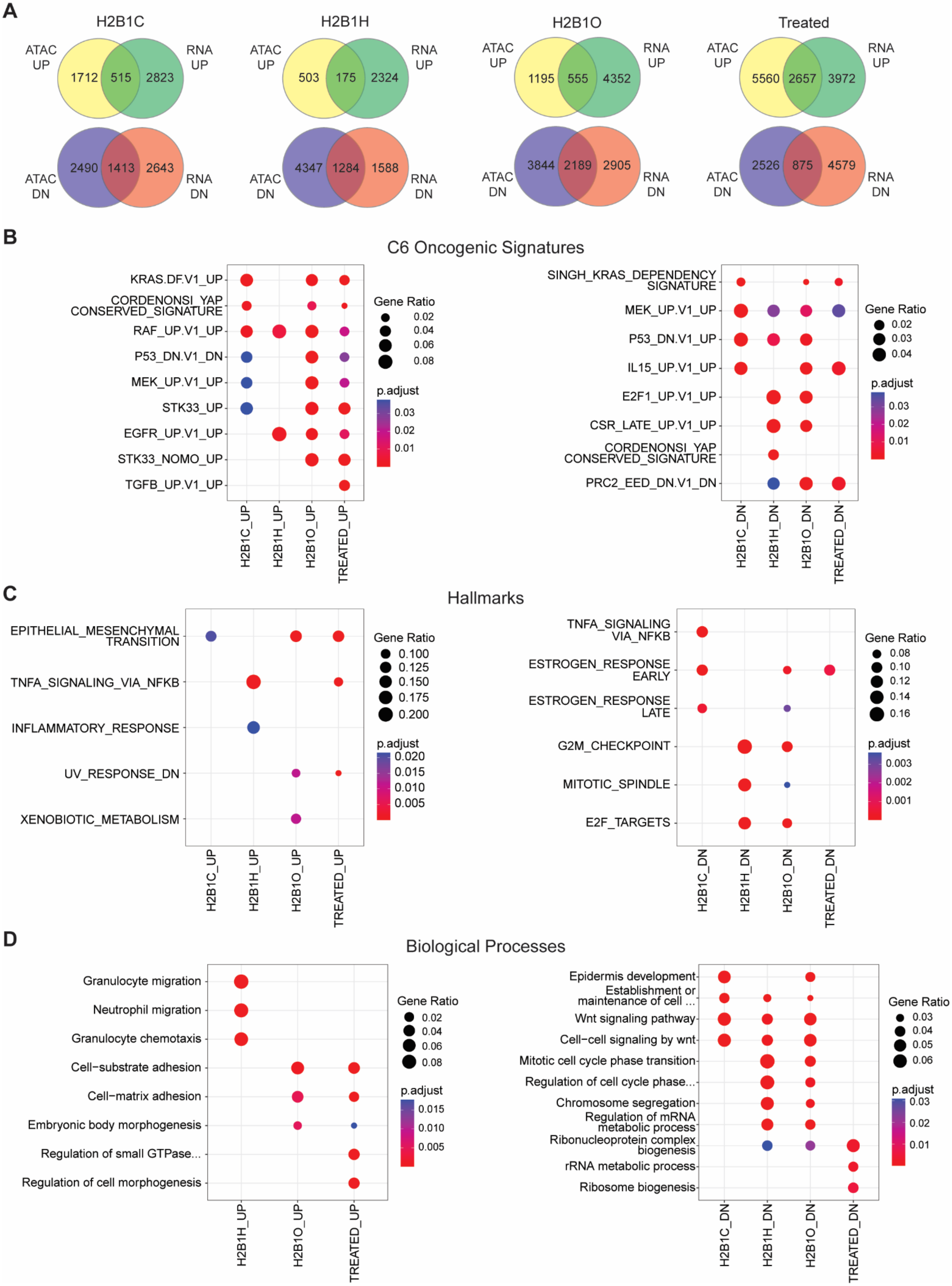
A
H2B1C
ATAC UP
RNA UP
1712
515
2823
2490
1413
2643
ATAC DN
RNA DN
H2B1H
503
175
2324
4347
1284
1588
H2B1O
1195
555
4352
3844
2189
2905
Treated
5560
2657
3972
2526
875
4579
B
C6 Oncogenic Signatures
KRAS.DF.V1_UP
CORDENONSI_YAP_CONSERVED_SIGNATURE
RAF_UP.V1_UP
P53_DN.V1_DN
MEK_UP.V1_UP
STK33_UP
EGFR_UP.V1_UP
STK33_NOMO_UP
TGFB_UP.V1_UP
H2B1C_UP
H2B1H_UP
H2B1O_UP
TREATED_UP
Gene Ratio
0.02
0.04
0.06
0.08
p.adjust
0.03
0.02
0.01
SINGH_KRAS_DEPENDENCY_SIGNATURE
MEK_UP.V1_UP
P53_DN.V1_UP
IL15_UP.V1_UP
E2F1_UP.V1_UP
CSR_LATE_UP.V1_UP
CORDENONSI_YAP_CONSERVED_SIGNATURE
PRC2_EED_DN.V1_DN
H2B1C_DN
H2B1H_DN
H2B1O_DN
TREATED_DN
Gene Ratio
0.02
0.03
0.04
p.adjust
0.03
0.02
0.01
C
Hallmarks
EPITHELIAL_MESENCHYMAL_TRANSITION
TNFA_SIGNALING_VIA_NFKB
INFLAMMATORY_RESPONSE
UV_RESPONSE_DN
XENOBIOTIC_METABOLISM
Gene Ratio
0.100
0.125
0.150
0.175
0.200
p.adjust
0.020
0.015
0.010
0.005
TNFA_SIGNALING_VIA_NFKB
ESTROGEN_RESPONSE_EARLY
ESTROGEN_RESPONSE_LATE
G2M_CHECKPOINT
MITOTIC_SPINDLE
E2F_TARGETS
Gene Ratio
0.08
0.10
0.12
0.14
0.16
p.adjust
0.003
0.002
0.001
D
Biological Processes
Granulocyte migration
Neutrophil migration
Granulocyte chemotaxis
Cell−substrate adhesion
Cell−matrix adhesion
Embryonic body morphogenesis
Regulation of small GTPase...
Regulation of cell morphogenesis
H2B1H_UP
H2B1O_UP
TREATED_UP
Gene Ratio
0.02
0.04
0.06
0.08
p.adjust
0.015
0.010
0.005
Epidermis development
Establishment or maintenance of cell ...
Wnt signaling pathway
Cell−cell signaling by wnt
Mitotic cell cycle phase transition
Regulation of cell cycle phase...
Chromosome segregation
Regulation of mRNA metabolic process
Ribonucleoprotein complex biogenesis
rRNA metabolic process
Ribosome biogenesis
H2B1C_DN
H2B1H_DN
H2B1O_DN
TREATED_DN
Gene Ratio
0.03
0.04
0.05
0.06
p.adjust
0.03
0.02
0.01

Fig S9, continued

E

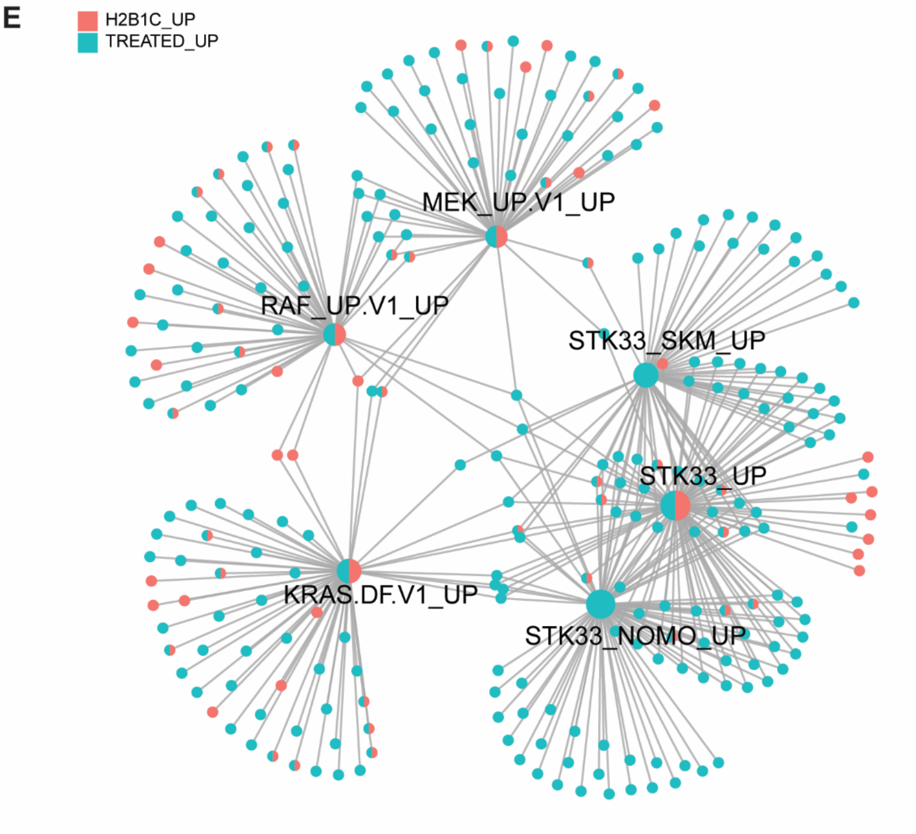


F

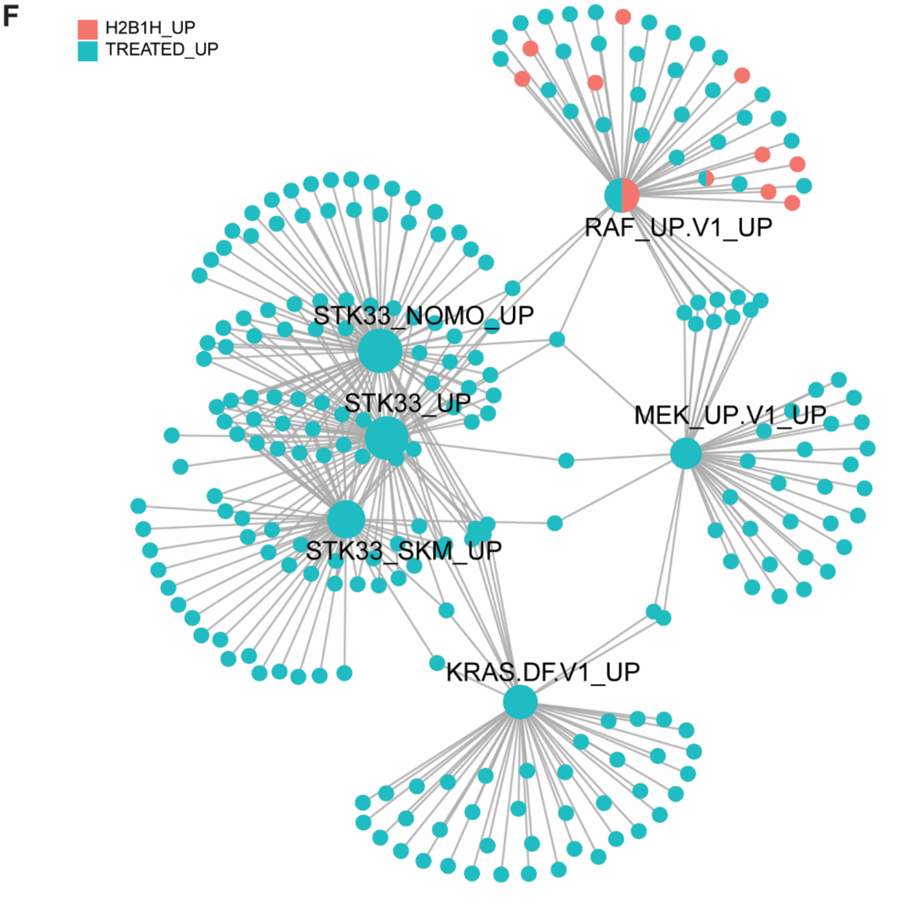

Fig S9, continued

**G**

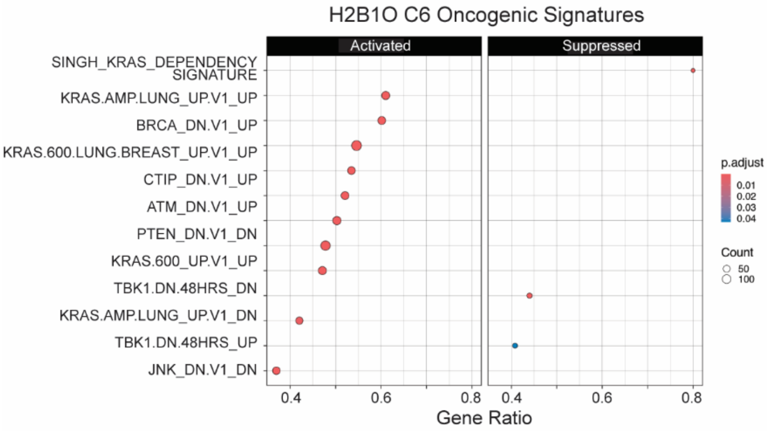


See Fig 8B for H2B1O Hallmarks

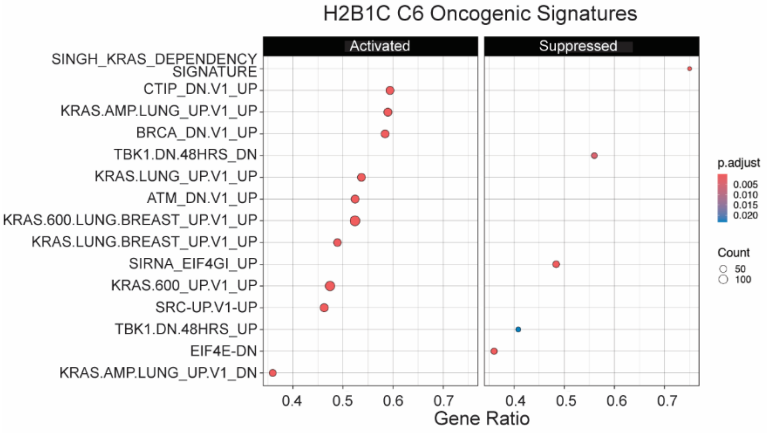


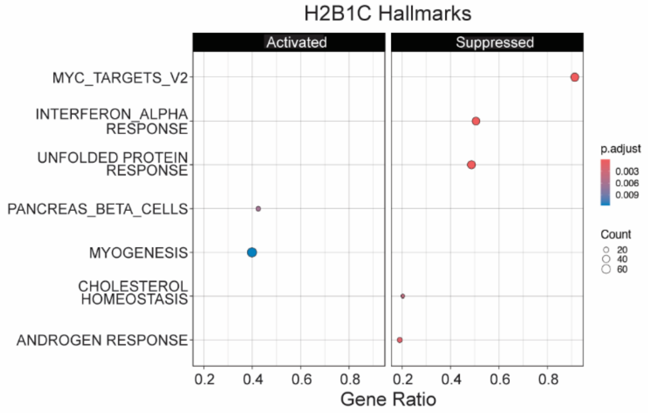


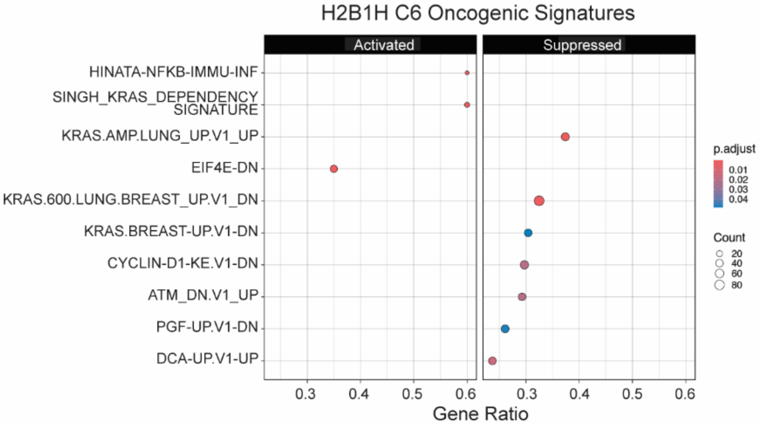


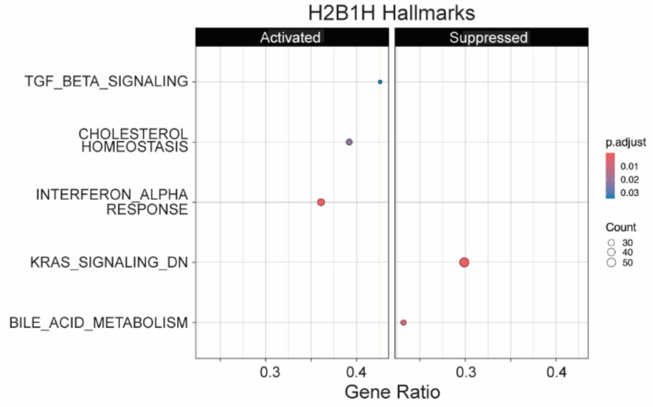
